# Odd-even disparity in the population of slipped hairpins in RNA repeat sequences with implications for phase separation

**DOI:** 10.1101/2023.01.09.523227

**Authors:** Hiranmay Maity, Hung T. Nguyen, Naoto Hori, D. Thirumalai

## Abstract

Low complexity nucleotide repeat sequences, which are implicated in several neurological disorders, undergo liquid-liquid phase separation (LLPS) provided the number of repeat units, *n*, exceeds a critical value. Here, we establish a link between the folding landscapes of the monomers of trinucleotide repeats and their propensity to self-associate. Simulations using a coarse-grained Self-Organized Polymer (SOP) model for (CAG)_*n*_ repeats in monovalent salt solutions reproduce experimentally measured melting temperatures, which are available only for small *n*. By extending the simulations to large *n*, we show that the free energy gap, Δ*G*_*S*_, between the ground state (GS) and slipped hairpin (SH) states is a predictor of aggregation propensity. The GS for even *n* is a perfect hairpin (PH) whereas it is a SH when *n* is odd. The value of Δ*G*_*S*_ (zero for odd *n*) is larger for even *n* than for odd *n*. As a result, the rate of dimer formation is slower in (CAG)_30_ relative to (CAG)_31_, thus linking Δ*G*_*S*_ to RNA-RNA association. The yield of the dimer decreases dramatically, compared to the wild type, in mutant sequences in which the population of the SH is decreases substantially. Association between RNA chains is preceded by a transition to the SH even if the GS is a PH. The finding that the excitation spectra, which depends on the exact sequence, *n*, and ionic conditions, is a predictor of self-association, should also hold for other RNAs (mRNA for example) that undergo LLPS.

## Introduction

The pioneering study by Eisenberg and Felsenfeld^1^ showed that polyriboadenylic acid (poly rA) could undergo phase separation in which a dense phase (condensate) coexists with the dispersed (sol) phase. However, it is only recently the importance of condensate formation involving RNA molecules in a variety of contexts is starting to be appreciated.^2–10^ The diversity of RNA sequences as well as the myriad structures an isolated RNA (a self-organizing polymer) adopts make it difficult to uncover the molecular features that drive phase separation. Nevertheless, one could anticipate a few scenarios for phase separation by favorable RNA-RNA interactions. (1) RNA molecules, containing unpaired single strand regions, may have a high propensity to associate with other RNA molecules, through complementary base pair formation, to form aggregates.^11,12^ In principle, both homotypic and heterotypic interactions are possible but in many cases one observes predominantly homotypic interactions, especially when there is sequence diversity.^4,7^ (2) The presence of a single strand in the ground state (lowest free energy state) that could engage in Watson-Crick (WC) base pair formation with other RNA chains is not always required. Even if the ground state is perfectly ordered, with no discernible disordered regions, phase separation does take place, as was shown in experiments^8,10^ and confirmed using simulations^13^ for repeat RNA sequences. (3) Because RNA is a polyanion, it stands to reason that cations could be efficacious in promoting phase separation by forming bridging interactions across RNA chains, although the mechanism of how this could be achieved is unknown.^14^

Regardless of the scenarios, the driving force for aggregate formation must involve favorable free energy of association in order to compensate for the entropy loss due to confinement of the chains in droplets that, besides restricting exploration of all allowed conformations, also inhibits both translational and rotational motions. The free energy of association leads to an increase in the number of intermolecular WC base pair formation per chain relative to the monomer in isolation as well as enhanced entropy^13,15^ in the increased ways in which the base pairs can be paired in a condensate containing many chains. These arguments raise the following question: Could one anticipate the propensity of RNA chains to self-associate using the properties of RNA in the monomeric form? We answer this question in the affirmative for low complexity repeat RNA sequences, although the findings maybe general.

Trinucleotide repeats, ubiquitous in human transcriptomes, are found in both the untranslated and translated regions of the human genome.^16^ A number of neurological and neuromuscular diseases, such as Huntington (HD), muscular dystrophy and amyotrophic lateral sclerosis (ALS), are related to the expansion of repeat sequences that exceed a critical length.^17^ For instance, in healthy humans, the repeat length of cytosine-adenine-guanosine (CAG) Huntington genes varies from 16 to 20. In contrast, the length of CAG, resulting in HD, exceeds a critical value (∼ 35).^18^ In an insightful paper, Jain and Vale (JV)^8^ showed that (CAG)_n_ polymers form biomolecular condensates provided *n* exceeds a critical number, which is close to the value required for the onset of the diseases. Here, we provide a quantitative description of the folding landscapes of (CAG)_n_ monomers, as a function of *n* in monovalent salt solutions, in order to determine the nature of states that drives self-association between RNA chains.

We developed a coarse-grained (CG) model that is sufficiently accurate to simulate (CAG)_n_ for arbitrary *n*. This genre of models has been proven to be useful in simulating the folding thermodynamics and kinetics in a variety of RNA molecules,^19–23^ with virtually no restriction on *n* or sequence diversity. Following our recent study,^13^ we used the Self-Organized Polymer (SOP) RNA model in which each nucleotide is represented by a single interaction site.^24^ Using the SOP model, we first investigated the structure and thermodynamics of (CAG)_n_ as a function of *n* and monovalent salt concentration. The model predicts the melting temperatures that are in good agreement with available experiments. The populations of hairpins with perfectly aligned strands (PH) and slipped hairpins (SH), containing one or more overhangs, depend on whether *n* is even or odd and the concentration of monovalent salt, *C*_*s*_. The ground state (GS) for odd (even) *n*, is the SH (PH). The slipped end in the SH is the source of multivalent interactions between the RNA polymers, which promote association of (CAG)_n_. The free energy gap, Δ*G*_*S*_, between the GS and the excited state with (at least) one slip is a predictor of the propensity to form dimers, and possibly the high density droplets. We validate this proposal by showing that the dimer formation rate in (CAG)_n_, with *n*=31, is greater than for *n*=30. Our work suggests that free Δ*G*_*S*_ between the GS and an excited state, with minimum of one CAG overhang, is a good indicator of the propensity for self-association.

## Results

### Calculated melting temperatures for small *n* agree with experiments

We first validated the SOP model simulations by comparing the calculated melting temperatures (*T*_*M*_s) with experiments. In the ground states of (CAG)_n_, nucleotides G and C form Watson-Crick base pair with three hydrogen bonds. The only unknown parameter in the SOP model is the hydrogen bond strength 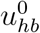 (see Eq. S3 in the Supplementary Information (SI)) whose value is determined, as described in the SI. We first performed folding simulations for the sequences, G(CAG)_n_C (*n* = 5, 6, and 7) at the monovalent salt concentration, *C*_*s*_ = 1 M. For these sequences, the experimental melting temperatures at *C*_*s*_ = 1 M^25^ have been measured, which we used to extract the value of 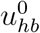 for use in all the simulations (see Fig. S1 and the SI for details).

For each 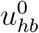, we performed simulations in the temperature range 0° ≤ *T* ≤ 127° C, and calculated the heat capacity, 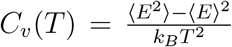 (where ⟨*E*⟩ is the average of the potential energy, ⟨*E*^2^⟩ is the associated mean square value), as a function of *T*. The location of the maximum in *C*_*v*_(*T*) is the melting temperature, *T*_*M*_. The calculated *T*_*M*_ values, with 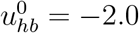 kcal/mol, are 49°C, 68°C, and 77°C for G(CAG)_5_C, G(CAG)_6_C and G(CAG)_7_C, respectively. The corresponding experimental values are ≈ 65°C, 68°C, and, 69°C, respectively. The fair agreement for *T*_*M*_s between experiments and simulations (Fig. 1 A) shows that the model, with a single parameter 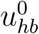, is a reasonable starting point for predicting the outcomes for longer repeat lengths. In the simulations of all other sequences, we fixed 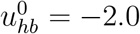 kcal/mol.

**Figure 1:**
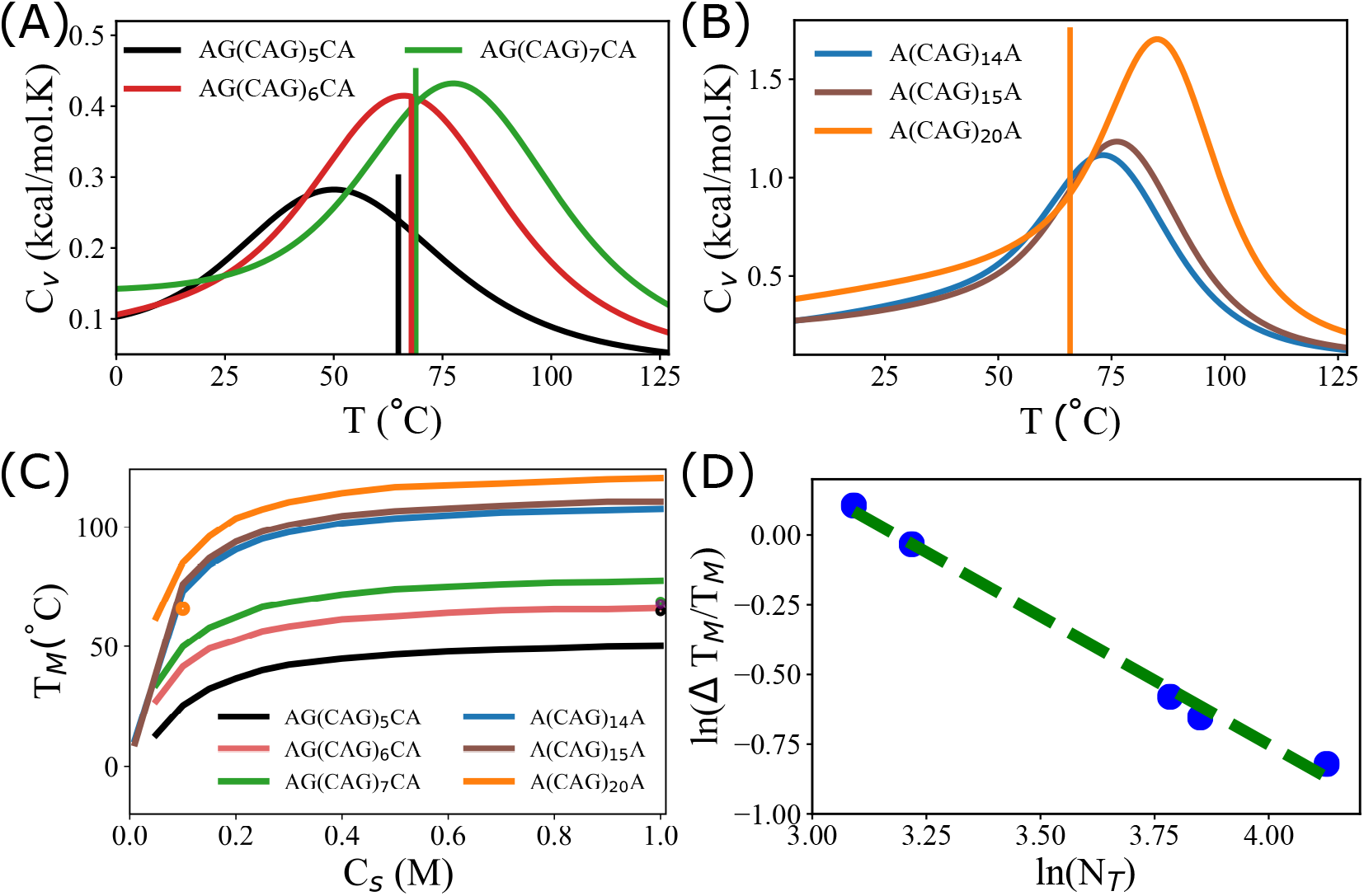
Thermodynamics of (CAG)_*n*_sequences: (A) Heat capacity, *C*_*v*_, as a function of temperature, *T*, for G(CAG)_5_C (black), G(CAG)_6_C (red), and G(CAG)_7_C (green) lines. The solid lines correspond to the experimental melting temperatures, *T*_*M*_s. (B) Same as (A) except the plots are for (CAG)_14_, (CAG)_15_ and (CAG)_20_ are in blue, cyan and magenta lines, respectively. Experimental value for *T*_*M*_ for (CAG)_20_ is shown as a solid line. (C) Predictions of *T*_*M*_s as a function of *C*_*s*_ for G(CAG)_n_C for various *n* that are indicated in the plot. The symbols are the measured values. (D) Log-Log plot of 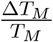 (Δ*T*_*M*_ is the full width at half maximum in *C*_*v*_(*T*)) as a function of the number of nucleotides, *N*_*T*_. The slope of the line is ≈ 0.92, which is close to the expected value of unity based on thermodynamic considerations.

### Melting temperatures of (CAG)_*n*_ as a function of salt concentration

We then simulated the folding of six (CAG)_*n*_ (with 5 ≤ *n* ≤ 20) as a function of *C*_*s*_. In addition to *n* = 5, 6 and 7 (Fig.1 A), we also calculated *C*_*v*_ as a function of temperature at *C*_*s*_ = 0.1 M for (CAG)_n_ with *n* = 14, 15 and 20 (Fig.1B). The calculated *T*_*M*_ (≈ 81° C) for (CAG)_20_ at *C*_*s*_ = 0.1 M is in reasonable agreement with the experimental value (≈ 66° C).^26^ In principle, better agreement could be obtained by adjusting 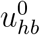. Because our goal is to find general principles that govern the propensity of RNA sequence to phase separate, we refrained from carrying out this exercise.

The *C*_*s*_-dependent melting temperatures change substantially up to *C*_*s*_ = 0.2 M. As *C*_*s*_ increases further, the *T*_*M*_ increase almost linearly up to *C*_*s*_ = 1 M (Fig.1C and Fig. S2 in the SI). The calculated melting temperatures for G(CAG)_5_C, G(CAG)_6_C, and G(CAG)_7_C agree well with the experimental values determined by UV-absorbance measurements done at 1M monovalent (Na^+^) salt concentration. The value of *T*_*M*_ for (CAG)_20_ at *C*_*s*_ = 0.1M obtained using simulations (*T*_*M*_ ≈ 86° C) differs by about 20° C (Fig. 1B) from experimental (*T*_*M*_ ≈ 66°C), as reported in Table 3 in^26^).

There could be two reasons for the discrepancy: (1) Experimental melting curves were measured using the standard UV absorbance^26^ whose relation to the theoretical calculations based on energy fluctuations is unclear. (2) It is possible that the Debye-Huckel approximation used in the simulations is not accurate at low Na^+^ or K^+^ concentrations.

### Finite size effects on *T*_*M*_

As *n* increases, the width of the heat capacity curves decreases, which indicates that the first-order transitions are sharper (Fig. 1A and Fig. 1B). From general thermodynamic considerations it can be shown that the ratio 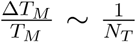 where *N*_*T*_ is the total number of nucleotides. Previously, we have shown that this relation holds for proteins.^27^ Fig. 1D shows that for the (CAG)_*n*_ the scaling holds perfectly.

### Populations of perfect hairpin (PH) and slipped hairpin (SH) vary for even and odd *n*

To understand the propensities to form condensates, we characterized the ensemble of structures explored by (CAG)_*n*_ as a function of *n*. The (CAG)_*n*_ sequences adopt multiple stem-loop or hairpin structures depending on the arrangement of the base pairs (bp) in the stem region. We characterized the hairpin structures using the order parameter, *Q*_*HP*_ (Eq. 1), which quantifies the deviations in the arrangement of the bps of a given structure from a perfectly aligned hairpin (PH) (Fig.2C and Fig.2D). An ensemble of conformations with *Q*_*HP*_ = 0 implies formation of bps that are identical to a PH with no mismatches, except the unavoidable A-A mismatch. The set of bps, *S*_*bp*_, in a given conformation can be expressed using, *S*_*bp*_ = (*i, j*) : *i* + *j* = *N*_*T*_ + 1, where, *i* and *j* are the indices of the nucleotides that form a base pair, and *N*_*T*_ is the total number of nucleotides in the sequence. In contrast to perfect hairpins, *Q*_*HP*_ = *m* (*m >* 0) where the positive integer *m* is a measure of strand slippage, which can occur either at the 5*′* or the 3*′* ends. Such structures represent deviations from PH.

**Figure 2:**
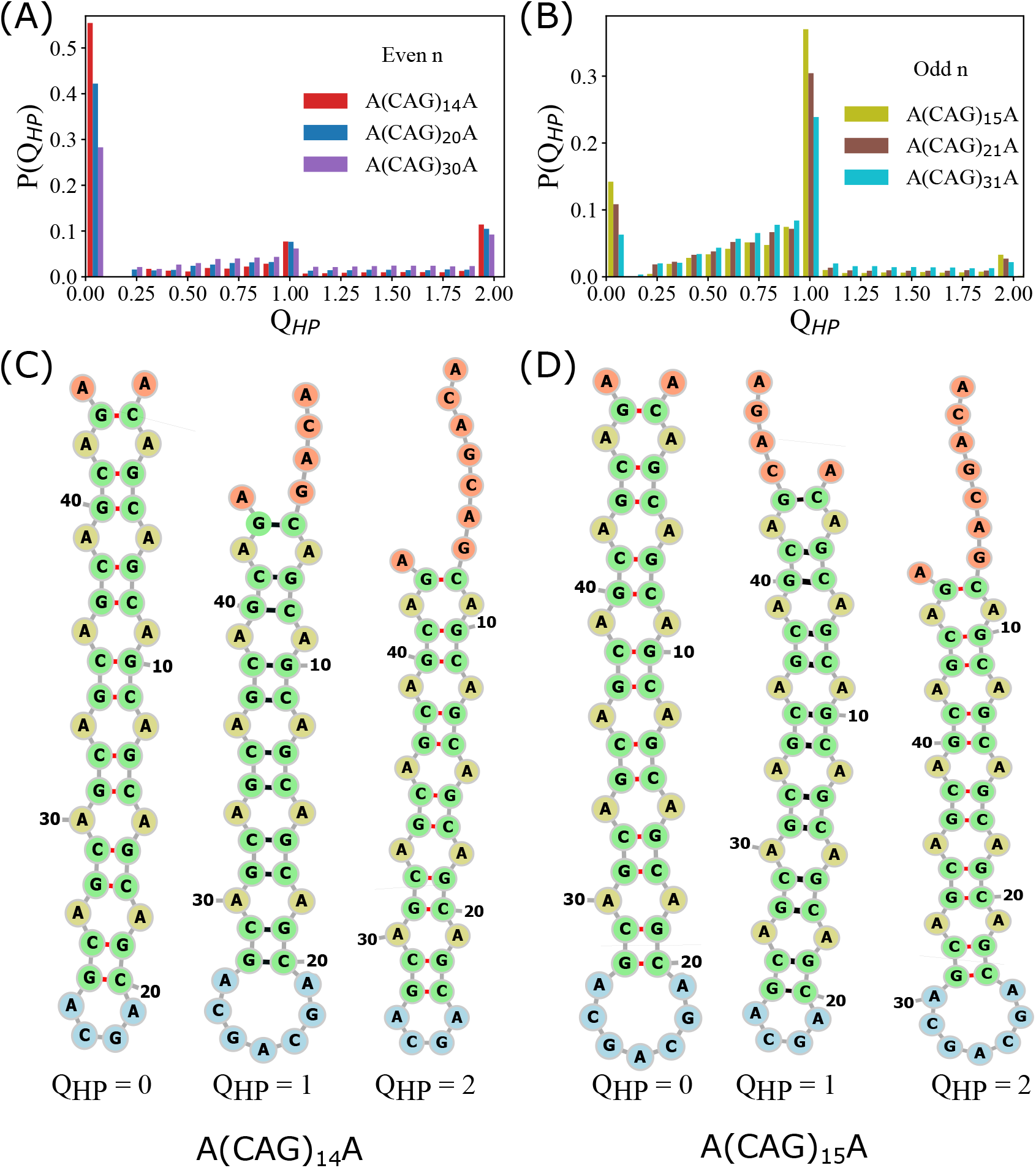
Characterizing the hairpin-like structures: (A) Distribution, *P*(*Q*_*HP*_), of the order parameter *Q*_*HP*_, for (CAG)_14_, (CAG)_20_ and (CAG)_30_. The maximum in *P*(*Q*_*HP*_) is at *Q*_*HP*_ = 0, implying that the ground state is a PH. (B) Same as (A), except *P*(*Q*_*HP*_)s are shown for odd values of *n*. The ground states have one unit of slippage, resulting in *Q*_*HP*_ = 1. (C) Representative hairpin structures with different *Q*_*HP*_ for (CAG)_14_ as an example. (D) The most populated states along with the *Q*_*HP*_ values for (CAG)_15_. Structures with fractional *Q*_*HP*_ are displayed in the SI.

Conformations, with fractional *Q*_*HP*_, typically have one or more bulges in the stem (Fig. S8 C) region. The classification using *Q*_*HP*_ allows us to quantitatively represent the folding landscape of repeat RNAs in terms of the ground and excited states of the chain. We show below that the free energy spectrum of states gives quantitative insights into the propensity for the RNA chains to self-associate.

We also calculated the probability distributions, *P*(*Q*_*HP*_)s, for the six sequences at *C*_*s*_ = 100 mM and temperature = 37°C (Fig. 2). The most populated structure (GS) for (CAG)_n_ depends on whether *n* is odd (*n* = 2*k* + 1) or even (*n* = 2*k*). For *n* = 2*k* the state with the highest population is a PH with *Q*_*HP*_ = 0 (Fig. 2A) whereas for *n* = 2*k* + 1 the value of *Q*_*HP*_ = 1 in the ground state (Fig. 2B). It is worth noting that the GS population decreases as *n* increases regardless of whether *n* is odd or even. Interestingly, for even *n*, the population of the excited state with *m* = 2 (two slippage units) is modestly higher than with *m* = 1. For both *m* = 1 and *m* = 2, the number of base pairs is the same in the stem. However, formation of the hairpin-like structure with *m* = 1 requires a loop with seven nucleotides whereas the excited state with *m* = 2 can be accommodated by the more stable tetraloop.

In addition to the GS, there are states with non-integer values of *Q*_*HP*_ that are also populated, albeit with smaller probabilities. The populations of such structures are greater if *n* is odd compared to even *n* (Figures 2A and B). The accessibility of such low free energy structures for (CAG)_31_ makes aggregation propensity greater than in (CAG)_30_ (see below).

Our findings that (CAG)_n_ with odd (even) number of repeats form hairpin with slippage (no slippage) in strands are in accord with experiments^28^ on single stranded (CAG)_n_ DNA and (CAG)_17_ RNA.^29^ The GS configuration obtained for odd or even numbered repeats is also consistent with the prediction from the RNAfold web server generated at *T* =37°C and *C*_*s*_ = 1 M. Experiments^30^ probing the mechanism for the conversion of (CTG)_*n*_ DNA hairpins between blunt end (*Q*_*HP*_ = 0) and overhang (similar to *Q*_*HP*_ = 1) configurations suggest the conversion occurs through the propagation of bulge (0 *< Q*_*HP*_ *<* 1) in the stem.

### Factors contributing to the stability of the ground states

Alternation in the ground state structure in going from *n* = 2*k* to 2*k* + 1 may be explained using the stability of both the loop and stem regions. Representative hairpin structures for even and odd numbered repeats are given in Figs. 2C and 2D, respectively (see also Fig. S8 in the SI). For *n* = 2*k*, the GS has *Q*_*HP*_ = 0 because such conformations accommodate the maximum number of bps, 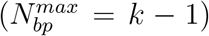 in the stem region. In addition to *Q*_*HP*_ = 0, the hairpins with *Q*_*HP*_ ≤ 1 also have the 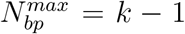 provided *n* = 2*k* + 1. As argued above, the preference for *Q*_*HP*_ = 1 over other structures is attributed to the stability of the loop region. All other conformations with 0 *< Q*_*HP*_ *<* 1 contain at least one bulge in the stem (see Fig. S8 in the SI for examples). Electrostatic repulsion generated between the negatively charged phosphates of the unpaired nucleotides is a minimum for *n* = 2*k* +1 if the sequence is in the *Q*_*HP*_ =1 state. Thus, loop entropy and electrostatic interactions determine the ground states of the low complexity sequences.

### (CAG)_31_ dimerizes faster than (CAG)_30_

Our previous work^13^ showed that homotypic interactions between (CAG)_*n*_ chains results in the formation of droplets that coexist with monomers (or small oligomers) if *n* is large enough (typically around 30). Examination of the structural changes that occur when a single polymer chain adds on to a droplet revealed that a slipped state (initially with *m* = 1) forms first either from the 5*′* or the 3*′* end, exposing unsatisfied CG bases. Subsequently, the CG bases engage in complimentary interactions with other chains in the droplet^13^. This picture suggests that the ease of formation of the SH states could reveal the propensity to form droplets or duplexes or low order oligomers. Therefore, we expect that the Δ*G*_*S*_ = *G*_*m*_−*G*_*GS*_, the difference in energy between the ground and the SH state, typically with one slippage (*m* = 1) at either end of the chain, should correlate with duplex formation. If the GS is SH, as is the case for (CAG)_31_, then Δ*G*_*S*_ = 0, which means that the GS is poised to interact with another RNA chain in order to initiate dimer formation. If this physical picture is correct, we expect that the time for forming a dimer should be smaller for (CAG)_31_ compared to (CAG)_30_ because Δ*G*_*S*_ for (CAG)_30_ is larger than for (CAG)_30_ (compare the spectra in Fig.3A and Fig.3B).

**Figure 3:**
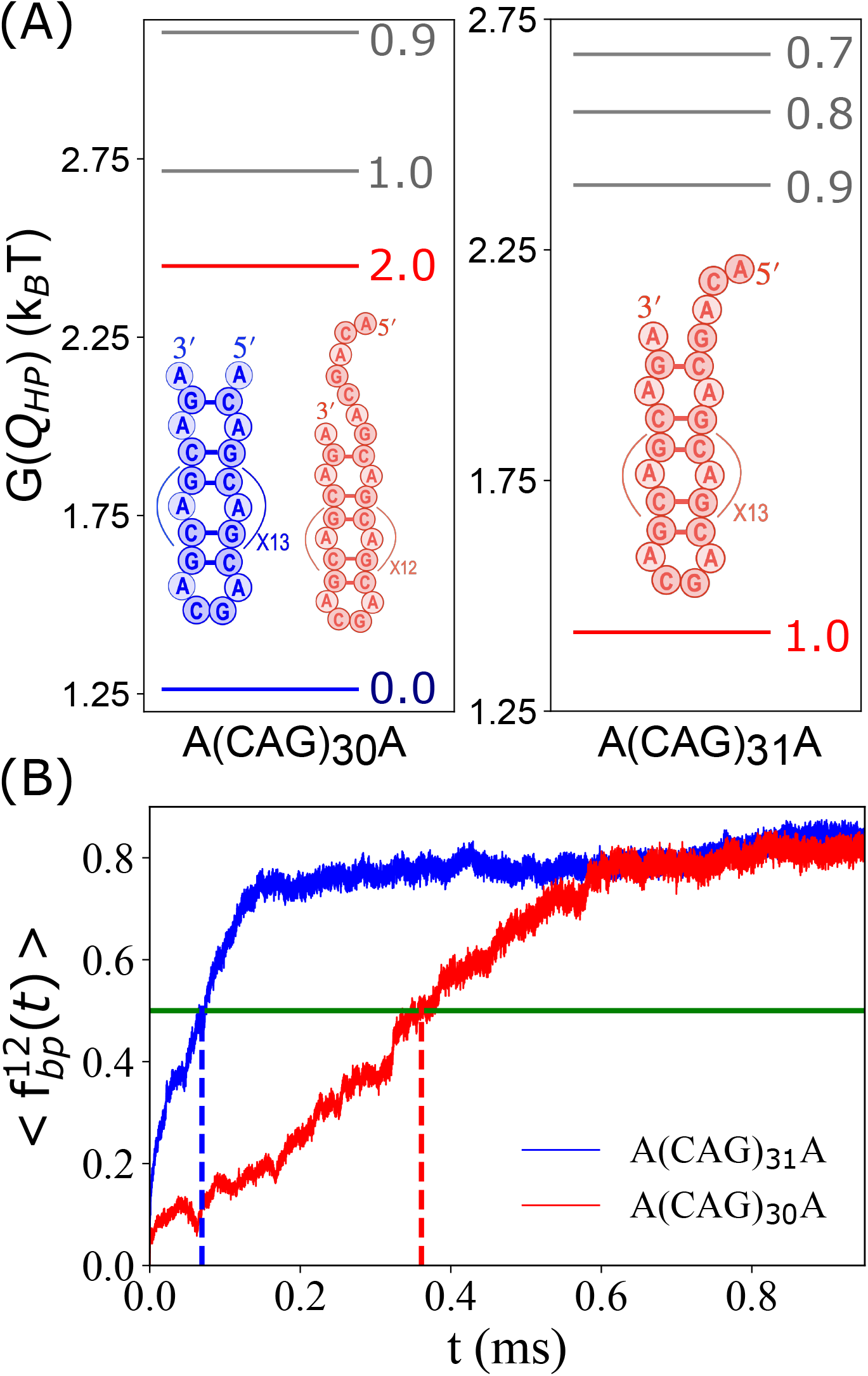
Link between free energy spectra and the kinetics of dimerization: (A) Free energy spectra for (CAG)_30_ and (CAG)_31_ computed at *C*_*s*_ = 0.1 M and *T* = 37° C. The value of *Q*_*HP*_ in the ground state is zero. Spectrum of states for (CAG)_31_, under the same conditions as in the left panel, is on the right. The ground state for odd *n* is slipped (*Q*_*HP*_ =1). (B) Time-dependent increase in the fraction of inter-chain base pairs, 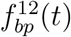 upon dimerization for (CAG)_30_ and (CAG)_31_. (CAG)_31_ dimer forms nearly 5 times faster than (CAG)_30_ dimer. Initially (*t* = 0) both the chains are in the ground state.

From the calculated spectra for these two constructs (Fig.3A and Fig.3B), we expect that (CAG)_31_ should dimerize faster than (CAG)_30_. To test this proposal, we simulated duplex formation (see the SI for details) for (CAG)_30_ and (CAG)_31_ by weakly constraining the polymers to remain within *R*_0_ = 40 Å(≈ ⟨*R*_*g*_⟩, the monomer radius of gyration). The dimer simulations were initiated from the monomer ground states which is a PH (SH) for (CAG)_30_ ((CAG)_31_). To monitor dimer formation, we calculated the fraction of inter molecular bp between the two chains 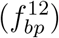 as a function of time. We assumed that a dimer is formed if 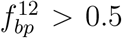. Out of the 100 trajectories, only in ≈ 40 % of the trajectories, the two chains adopted an anti parallel duplex (double stranded RNA) structure. Figure 3C shows the fraction, 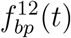, as a function of *t*. Based on the criterion that a dimer is formed if 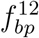 exceeds 0.5, we find that dimer formation is about 5 times slower in (CAG)_30_ compared to (CAG)_31_, thus qualitatively affirming the expectation based on the free energy spectra in Fig. 3. The estimated time scales for duplex formation is a lower bound because the friction coefficient used in the simulations is much smaller than the value should be in water. Nevertheless, we believe that there ought to be link between the accessibility of slipped states to self-association kinetics should be fairly general.

### Dimerization occurs predominantly from the SH states

The conversion of hairpins to a dimer for (CAG)_31_ occurs by two major pathways. Details of the structural transformations that occur along the predominant assembly pathway (Path I) are shown in Fig.4 and Fig. S11 in the SI for the minor pathway (Path II). In both the pathways, the hairpin transitions between different conformations before successful dimerization. The obligatory state prior to dimerization is the formation of the slipped state, either at the 5*′* or 3*′* terminus. Dimer formation occurs in 35 out of 40 trajectories (Path I). In all these trajectories, the slipped end of one of the hairpin starts interacting with the slipped end of the other chain. The structural transitions, measured using the fraction of intra and inter chain base pairs as a function of time for one trajectory, show that the fraction of bp loss in each chain is compensated by gain in the inter-chain base pairs (Fig.4A). The transition to the duplex state occurs over a short time window (given by the rectangle in black in Fig.4A) during which bulk of the inter-chain base pairs 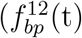 forms. Based on the mechanisms for the formation of dimers, we conclude that the propensity for the association of hairpins is higher if there is a slippage in the strand either at 5*′* or 3*′* termini. In other words, the free energy gap Δ*G*_*s*_, which is zero in (CAG)_31_ is the determining factor in dimer formation. In the minority path II (5 out of 40 trajectories), the slipped end of one hairpin interacts with the loop region of the other hairpin, and eventually forms the duplex structure (see the SI for details).

**Figure 4:**
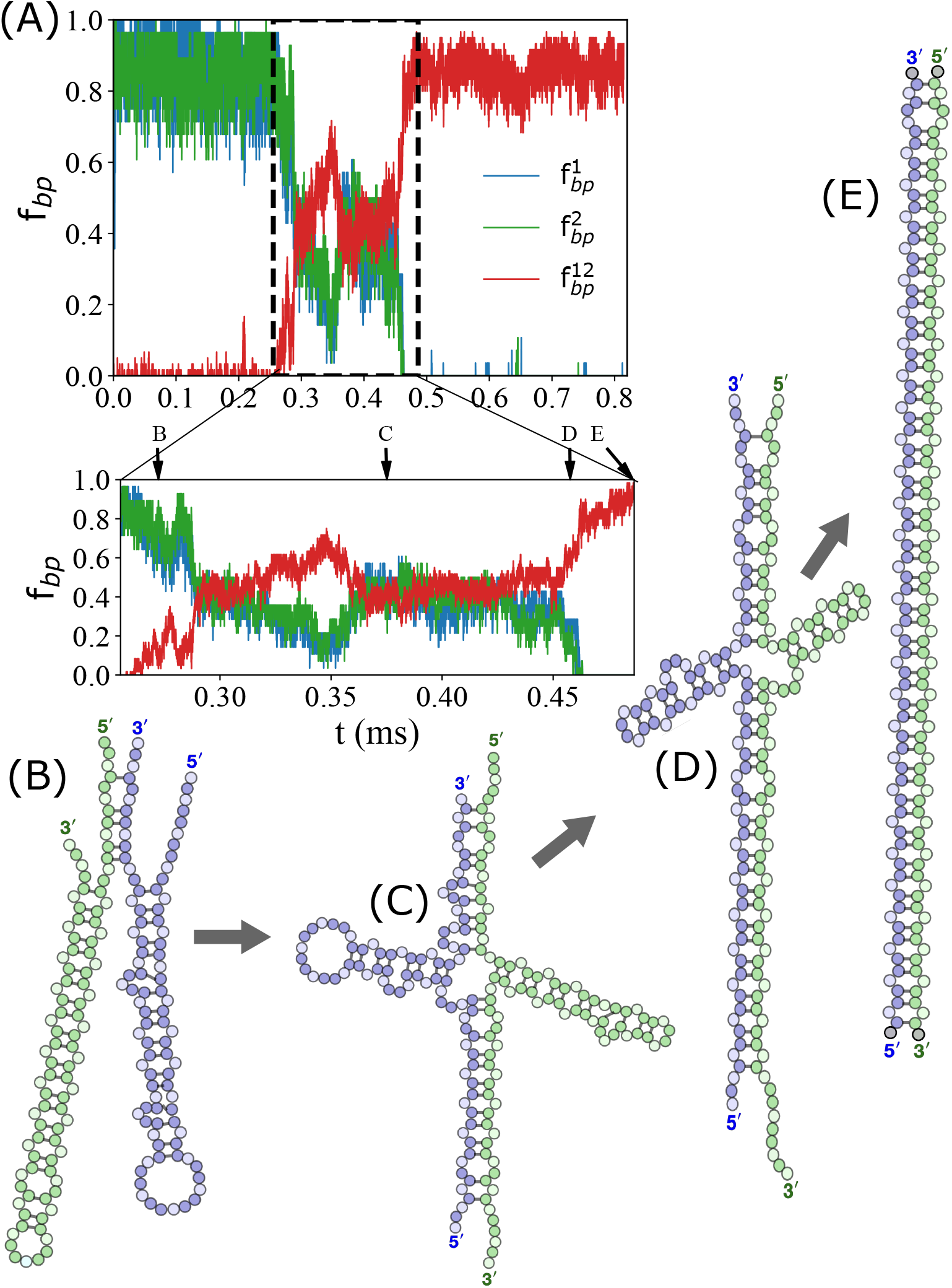
Major dimerization pathway formation for (CAG)_31_: (A) Time-dependent changes showing loss of fractions of intra-chain base pairs 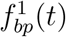 and 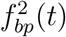 and gain in inter-chain base pairs 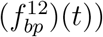 in the dominant pathway. The initial structures for both the chains correspond to their ground states (*Q*_*HP*_ = 1). The values of *C*_*s*_ is 100mM and *T* =37°. The panel below zooms in on the time range where the dimer forms. (B - D) Initial hairpin conformations first transition to the structures with a slipped ends at the termini. The hairpins with the slipped end start interacting with each other and eventually lead to the formation of a dimer through a sequence of transitions. Representative structures along the pathways are shown. Structural transitions in the minor pathway (Fig. S11) are shown in the SI.

The structural transitions that occur in Path I are sketched in Fig. 4B, C, D and E. They show that increases in the slippage in one chain result in the loss of intra chain WC base pairs. But the loss is compensated by an increase in the inter-chain base pairs, which eventually leads to the perfect duplex formation (Fig.4E). We showed in an earlier study^13^ that a similar mechanism holds in condensate formation as well. The sequence of transitions that occurs in a single RNA chain (see Fig. 6 in our previous study^13^) as it merges with the droplet is essentially the same as shown in Fig. 4. Thus, the driving force for condensate formation is the enhanced gain in forming not only many inter-chain CG base pairs but also the increase in the number of ways in which such base pairs form, which is an entropic factor.

### Effects of mutations on hairpin structures and dimer formation

A corollary of the predicted inverse correlation Δ*G*_*S*_ and the propensity to dimerize is that suppression in the population of SH should reduce the yield of the dimer. To test this prediction, we probed the effects of mutations on the population of hairpin structures using variants of (CAG)_30_ and (CAG)_31_. The M1 mutant sequence is A (CCG)(CAG)_28_(CGG)A, and the M2 sequence is A (CCG)(CAG)_29_(CGG)A. We also considered other sequences (see SI) but here we focus on M1 and M2.

We calculated the populations (*P*(*Q*_*HP*_) - shown in Fig. S12 in the SI) of various hairpin-like structures at *C*_*s*_ = 100 mM and *T* =37°C for M1 and M2 from which the free energy spectra is readily calculated (Fig.5A and B). The ground state of M1 is a PH with *Q*_*HP*_ = 0 whereas the GS of M2 is a structure with a bulge but has no slipped CAG even though *Q*_*HP*_ = 0.9 is close to unity (Fig.5). There are significant differences in *P*(*Q*_*HP*_)s between the WT and the mutants (Figure2 and S12). For example, *P*(*Q*_*HP*_ = 0) state in M1 increases significantly compared to the WT. In addition, Δ*G*_*S*_ for M1 is greater than in the WT. Comparison between the free energy spectra for the WT and M2 also shows dramatic differences (compare Fig.3A and Fig. 5B). The GS for the WT is a SH hairpin that is poised to form higher order structures by interacting with other RNA chains. The values of Δ*G*_*S*_ in the WT and M2 are smaller compared with Δ*G*_*S*_ between the WT and M1. From the results in Fig.4 and Fig.5, we expect that the yield of the dimer in mutant sequences should be considerably less than in the WT sequences.

**Figure 5:**
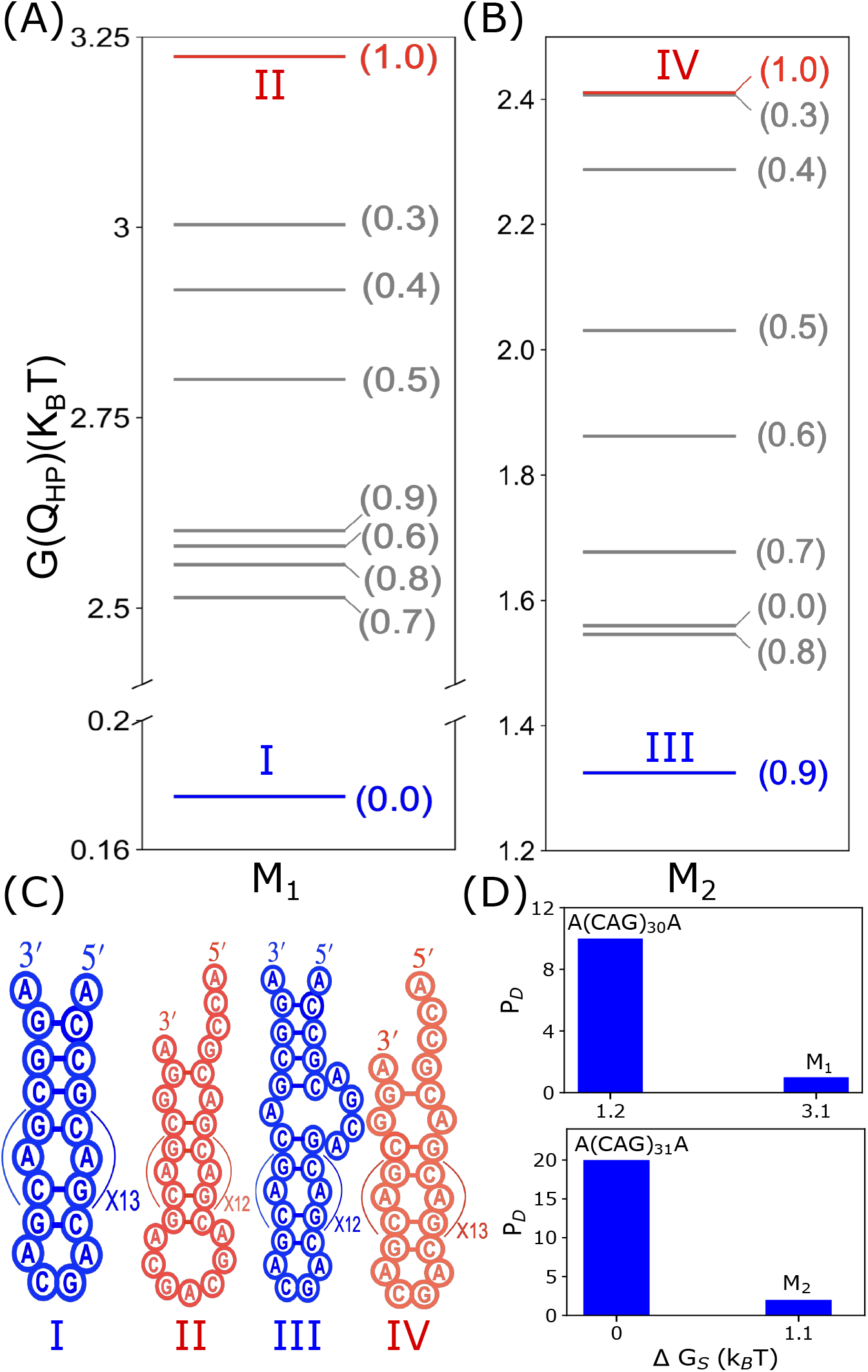
Effect of mutations on dimer yields: (A) Free energy spectrum of M1 mutant (A(CCG)(CAG)_28_(CGG)A). The numbers represent the values of *Q*_*HP*_. (B) Same as (A) except the folding landscape is for M2 (A(CCG)(CAG)_29_(CGG)A). The free energy between the ground state (PH) and the SH in M1 is higher than in M2. (C) Schematic structures of the relevant states (I, II, III, and IV) involved in dimer formation. (D) Dimer yields, expressed as percentage, for A(CAG)_30_A and M1 mutant are in the top panel, and the bottom panels show A(CAG)_31_A and M2. Note that the WT yield is higher in A(CAG)_31_A than in A(CAG)_30_A.

To illustrate the prediction that Δ*G*_*S*_ is the primary factor in determining the kinetics and yield of the duplex structures, we performed dimer simulations starting from the GS conformations for M1 and M2. Because the simulations are time consuming, we calculated the yield, instead of the rate of dimer formation. The percentage yield is calculated using, 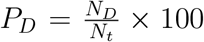 where *N*_*D*_ is the number of trajectories that reached the dimer state, and *N*_*t*_ is the total number of trajectories. We expect *P*_*D*_ to decrease as Δ*G*_*S*_ increases, which implies that *P*_*D*_ for the mutants must be greatly reduced relative to the corresponding WT sequences. There is a decrease in *P*_*D*_, by at least by one order of magnitude, in the mutant sequence compared to the WT sequences (Figure 5 C and D).

## Discussion

We created a coarse-grained model that allows for reasonably accurate calculations of the thermodynamic properties for monomer repeat RNA sequences as a function of monovalent salt concentration and sequence length. Because a minimum length of repeat sequences is required for condensate formation, we also simulated (CAG)_*n*_ for large *n* for both the wild type and several mutants, with the goal of linking their folding landscapes to the propensity of RNA polymers to self-associate. Two major findings that are likely to be valid for arbitrary RNA sequences emerge from our study. First, self-association is preceded by the formation of slipped hairpins even when most of the nucleotides are engaged in base pair formation. This is certainly the case when *n* is even. Second, there is an inverse correlation between the free energy gap separating the ground state and the aggregation prone slipped hairpin state, and the rates and yields of dimer formation. In other words, structured RNA, with no exposed single strand, is unlikely to phase separate if the excited state is inaccessible.

### Salt-dependence of folding thermodynamics

The finding that the melting temperature, *T*_*M*_, increases nearly linearly at small monovalent salt concentration, *C*_*s*_, and more slowly beyond *C*_*s*_ = 0.2M (Fig.1C and Fig. S2 in the SI) is amenable to experimental test. The free energy, ΔΔ*G*(*C*_*s*_) = Δ*G*(*C*_*s*_) − Δ*G*(0.15*M*), decreases as ΔΔ*G*(*C*_*s*_) ∼ −*kln*(*C*_*s*_) for all (CAG)_*n*_ (see Fig. S7 in the SI).

### Free energy spectra as a function of *n*

The differences between the ground state and the slipped hairpin free energies qualitatively explain the speedup in the dimer formation in (CAG)_31_ compared to (CAG)_30_. More precisely, it is the population of the SH that determines the aggregation rate. As *n* increases, we expect that Δ*G*_*S*_should decrease, which means that for large *n* the differences in the rates of self-association between RNA chains with odd and even *n* should decrease. A corollary of our finding is that mutations, which increase Δ*G*_*S*_should decrease dimerization rates, as illustrated in Fig. S15 in the SI. Finally, for a fixed *n* and identical sequence composition, the approximate dimer formation rate maybe altered by changing the sequence. Thus, the *exact* sequence,^10^ in addition to *n*, salt concentration and other environmental factors, determine the tendency to self-associate.

### Formation of a slipped hairpin is obligatory for self-association

The ground state of (CAG)_30_ is a perfect hairpin, and yet it self associates with another RNA chain to form a dimer. In order for this process to occur, both the RNA chains must transition to an excited state, resulting in the formation of SH, thus exposing one or more overhangs. The complimentary sequences then could hybridize, which would nucleate the formation of the dimer. Our previous study showed that monomer addition to a droplet involves initial transition to a slipped state (see Fig. 6 in^13^) that is followed by further unfolding that creates a large single stranded conformation. The resulting excited states can form complimentary base pairs with other chains in the droplet. Taken together, we ascertain that SH state formation, regardless of the ground state, is a necessary condition for RNA chains to self-associate. A corollary is that if the formation of the SH is prevented then droplets may not form or the time for self-association would greatly increase. This is the case in mutant sequences in which either the dimerization time increases or the yield of the dimer is diminished substantially.

**Figure 6:**
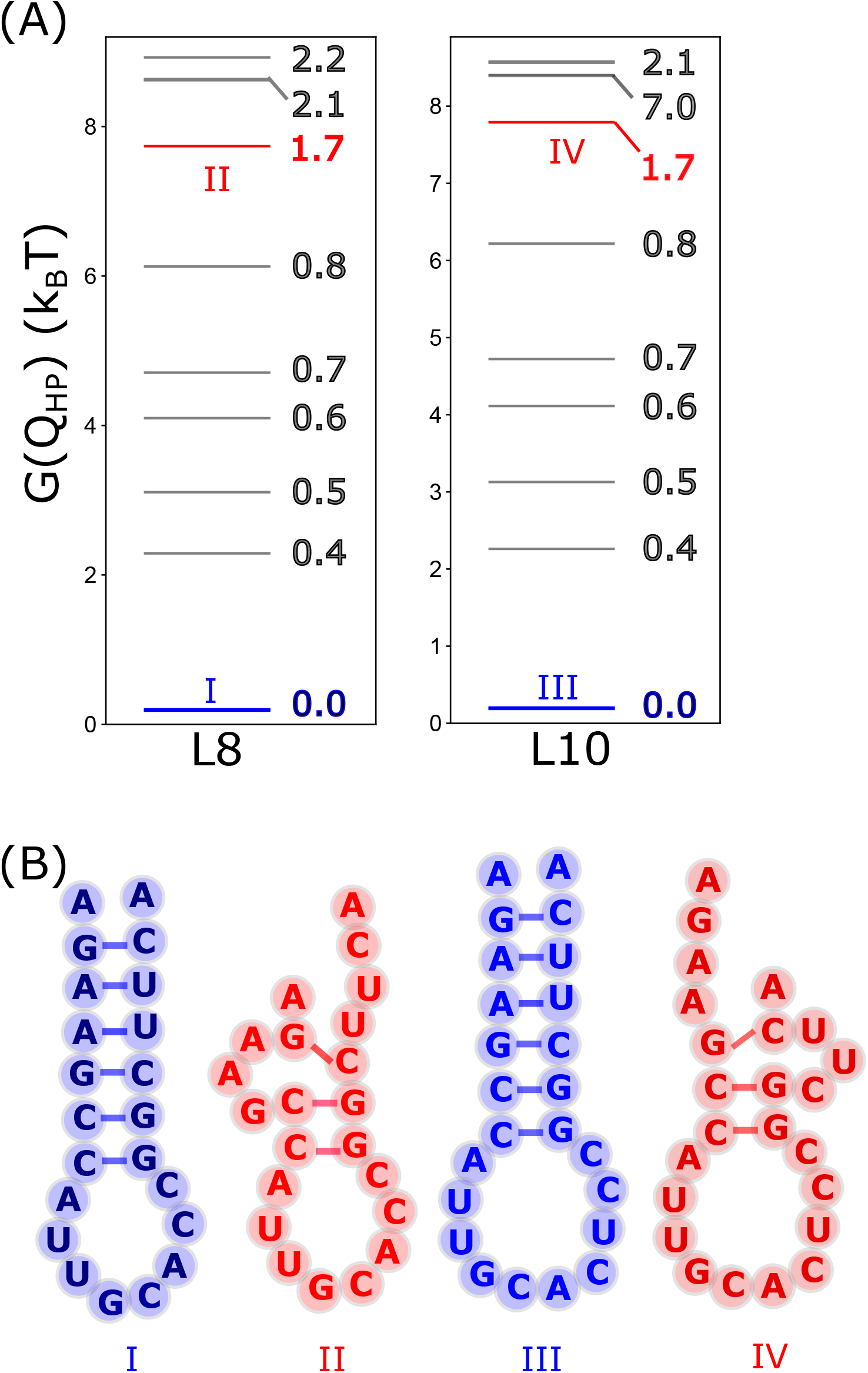
Free energy spectra and relevant structures for L8 and L10 hairpins: (A) Calculated spectra for hairpins with heterogeneity in the sequences. The free energies of the aggregation-prone states (in red) are separated from the ground states by about 8*k*_*B*_*T*, making the populations of such states vanishingly small. States with *Q*_*HP*_ *<* 1.7 do not have any slippage. (B) The ground state structures with 5*′* and 3*′* ends in proximity is in blue. The high free energy states, with slipped nucleotides, are in red.

### Sequence diversity should inhibit RNA-RNA interactions

Our findings raise a broader question: How are unwarranted interactions between various RNA chains, especially those that have to fold for functional reasons, in cells prevented? There are two possible reasons. (1) Self-association between RNA chains requires exposure of single stranded regions. Our findings suggest that, at least one of the two RNA chains, should be in the SH. Because Δ*G*_*S*_*/k*_*B*_*T* is not that large, the SH states are readily populated in low complexity sequences. A measure of complexity is the sequence entropy given by 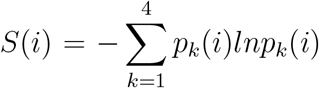 where *p*_*k*_(*i*) is the probability of finding nucleotide of type *k* (A,T,G,C) at position *i* in a multiple sequence alignment sense. For (CAG)_*n*_, *S*(*i*) ≡ 0 for all *i*. In more complex RNA sequences (*S*(*i*) *>* 0 - the maximum value is *ln*4), it is likely that Δ*G*_*S*_ is sufficiently large (exceeds ≈ 2*k*_*B*_*T*) that the population of states with exposed single strands is small. In order to demonstrate that this is indeed the case, we calculated the free energy spectra for two hairpins, L8 and L10 (Fig.6). The structure (labelled II in Fig.6B) with modest slippage (*Q*_*HP*_ =1.7) is separated from the ground state by ≈ 8*k*_*B*_*T* in L8 (left panel in Fig.6A). In L10, the first appearance of a slipped state (labelled IV in Fig.6B) in the free energy spectrum (Fig.6A) has *Q*_*HP*_ = 1.7 with a similar excitation free energy as in L8. Fig. S15 in the SI shows that in conformations with *Q*_*HP*_ *<* 1.7, the 5*′* and 3′ are close to each other. Therefore, their propensities of such structures to aggregate are diminished.

(2) RNA molecules, whose folding is needed for splicing (*Tetrahymena* ribozyme for example), misfold readily.^31–34^ However, the bases in the exposed regions in such misfolded states are often paired, although the structures are non-native containing bulges, loops, and multiloops. Although it is known that in the ground states of large single stranded RNA molecules the 5*′* and 3′ are in proximity,^35^ much less is known about the structures of the excited states. It is more than likely (L8 and L10 being very simple examples) that the exposed single stranded regions are not readily accessible in heterogeneous RNA sequences, which greatly suppresses the probability of association with other RNA molecules. Indeed, the excited state of P5abc sequence not only has low population (3%, which translates to Δ*G*_*S*_ ≈ 3.5*k*_*B*_*T*) but the NMR structure (Fig. 3 in^36^) shows absence of exposed single stranded regions. Therefore, we surmise that in P5abc, conformations with any slippage, that could form complimentary inter RNA interactions, must far exceed ≈ 3.5*k*_*B*_*T*, which implies that self-association of P5abc chains is unlikely.

## Conclusion

That the population of excited states determine protein aggregation propensity has been emphasized for a long time.^37,38^ Conceptually there are similarities between the monomer characteristics of protein monomers, and their tendencies to aggregate. Indeed, this analogy has been used to explain RNA-RNA interactions, especially when ribosome detaches from mRNA.^39^ The analogy to protein aggregation is most apt if the ground state of the protein is folded or native-like, which is the case in the amyloid formation in transthyretin and the associated mutants and aggregation of prion proteins.^40,41^ In both these examples, aggregation must involve access to partially misfolded states,^40,42,43^ which would expose hydrophobic residues, and facilitate protein aggregation. In contrast, amyloid formation in peptides (A*β*_42_ or Fused in Sarcoma) aggregation prone structures, with fibril like character, are the excited states while the ground states are disordered. Nevertheless, understanding self-association of RNA and proteins requires characterizing the entire free energy landscape, and not merely the ground states. Therefore, there is a great need for experimentally determining the free energies and structures of the excited states of RNA molecules, especially those that are involved in stress granule formation. This has been accomplished for proteins,^44–46^ and more recently for the P5abc domain.^36^

## Methods

We represent each nucleotide by a single site.^13,24^ To account for ion condensation onto the highly charged RNA polyanion, we used a reduced value of −*Q* on the phosphate groups with 0 *< Q <* 1.^47,48^ Oosawa-Manning counterion condensation theory^49,50^ was used to calculate the value of *Q*. The total energy, *E*_*T OT*_, of an RNA is the sum of bonded (*E*_*B*_), hydrogen bonded (*E*_*HB*_), excluded volume (*E*_*EV*_), and electrostatic energy (*E*_*el*_). The details of the model and simulations are given in the Supporting Information (SI)

### Structural Classification

In order to reveal the spectrum of free energy states of the RNA sequences, we first calculated, *P*(*Q*_*HP*_), the distribution of *Q*_*HP*_, where *Q*_*HP*_ is an order parameter that measures the deviation in the arrangement of base pairs with respect to a perfectly aligned hairpin structure. We define *Q*_*HP*_ as,

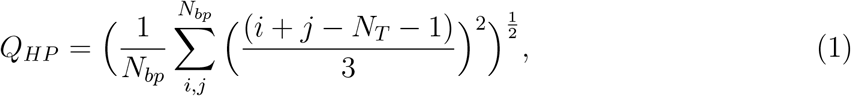

where, *i* and *j* label the nucleotides, *N*_*T*_ is the length of the sequence, and *N*_*bp*_ = number of base pairs in a given conformation. The nucleotide indices *i* and *j*, forming a base pair in a perfectly aligned hairpin, is related by *i* + *j* = *N*_*T*_ + 1. A hairpin with *Q*_*HP*_ = 0 shows that the strands are perfectly aligned with respect to each other (see the structures in Fig. 2C). Similarly, *Q*_*HP*_ = 1 corresponds to a slipped hairpin (SH) state (Fig. 2D). In the range 0 *< Q*_*HP*_ *<* 1 the conformations contains bulges in the stem (see Fig. S8 in the SI).

### Calculation of the free energy spectrum

In order to construct the free energy spectrum, we first arranged the population of different hairpin structures, *P*(*Q*_*HP*_), in descending order. The free energy of a given state, characterized by *Q*_*HP*_, is calculated using,

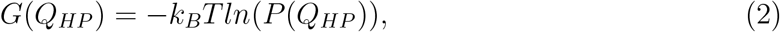

where, *k*_*B*_ is the Boltzmann constant and T is the absolute temperature. We constructed the free energy spectrum by considering the structures for which 0 ⪅ *G*(*Q*_*HP*_) ⪅ 4*k*_*B*_*T*. The structures with *G*(*Q*_*HP*_) *>* 4*k*_*B*_*T* are not shown because their populations are negligible.

## Acknowledgements

This work was supported by a grant from the National Science Foundation (CHE 19-00033) with additional support from the Welch Foundation (F-0019) through the Collie-Welch Chair.

## References

(1) Eisenberg, H.; Felsenfels, G. Studies of Temperature-dependent Conformation and Phase Separation of Polyriboadenylic Acid Solution at Neutral pH. J. Mol. Biol. 1967, 30, 17+.

(2) Roden, C.; Gladfelter, A. S. RNA contributions to the form and function of biomolecular condensates. Nat. Rev. Mol. Cell Biol. 2021, 22, 183–195.

(3) Van Treeck, B.; Protter, D. S. W.; Matheny, T.; Khong, A.; Link, C. D.; Parker, R. RNA self-assembly contributes to stress granule formation and defining the stress granule transcriptome. Proc. Natl. Acad. Sci. U. S. A. 2018, 115, 2734–2739.

(4) Trcek, T.; Douglas, T. E.; Grosch, M.; Yin, Y.; Eagle, W. V. I; Gavis, E. R.; Shroff, H.; Rothenberg, E.; Lehmann, R. Sequence-Independent Self-Assembly of Germ Granule mRNAs into Homotypic Clusters. Mol. Cell 2020, 78, 941+.

(5) Van Treeck, B.; Parker, R. Emerging Roles for Intermolecular RNA-RNA Interactions in RNP Assemblies. Cell 2018, 174, 791–802.

(6) Aumiller, W. M., Jr.; Cakmak, F. P.; Davis, B. W.; Keating, C. D. RNA-Based Coacervates as a Model for Membraneless Organelles: Formation, Properties, and Interfacial Liposome Assembly. Langmuir 2016, 32, 10042–10053.

(7) Trcek, T.; Grosch, M.; York, A.; Shroff, H.; Lionnet, T.; Lehmann, R. Drosophila germ granules are structured and contain homotypic mRNA clusters. Nat. Commun. 2015, 6.

(8) Jain, A.; Vale, R. D. RNA phase transitions in repeat expansion disorders. Nature 2017, 546, 243+.

(9) Langdon, E. M.; Qiu, Y.; Niaki, A. G.; McLaughlin, G. A.; Weidmann, C. A.; Gerbich, T. M.; Smith, J. A.; Crutchley, J. M.; Termini, C. M.; Weeks, K. M.; Myong, S.; Gladfelter, A. S. mRNA structure determines specificity of a polyQ-driven phase separation. Science 2018, 360, 922–927.

(10) Isiktas, A. U.; Eshov, A.; Yang, S.; Guo, J. U. Systematic generation and imaging of tandem repeats reveal base-pairing properties that promote RNA aggregation. Cell Reports Methods 2022, 2, 100334.

(11) Ma, W.; Zheng, G.; Xie, W.; Mayr, C. In vivo reconstitution finds multivalent RNA-RNA interactions as drivers of meshlike condensates. eLife 2021, 10.

(12) Langdon, E. M.; Gladfelter, A. S. A New Lens for RNA Localization: Liquid-Liquid Phase Separation; Annual Review of Microbiology; 2018; Vol. 72; pp 255–271.

(13) Nguyen, H. T.; Hori, N.; d, T. Condensates in RNA Repeat Sequences are Heterogeneously Organized and Exhibit Reptation Dynamics. Nature Chemistry 2022, p,.

(14) Onuchic, P. L.; Minn, A. N.; Alshareedah, I.; Deniz, A. A.; Banerjee, P. R. Divalent cations can control a switch-like behavior in heterotypic and homotypic RNA coacervates. Sci Rep 2019, 9.

(15) Kimchi, O.; King, E. M.; Brenner, M. P. Uncovering the mechanism for aggregation in repeat expanded RNA reveals a reentrant transition. bioRxiv 2022,

(16) Krzyzosiak, W. J.; Sobczak, K.; Wojciechowska, M.; Fiszer, A.; Mykowska, A.; Kozlowski, P. Triplet repeat RNA structure and its role as pathogenic agent and therapeutic target. Nucleic Acids Res. 2012, 40, 11–26.

(17) Everett, C.; Wood, N. Trinucleotide repeats and neurodegenerative disease. Brain 2004, 127, 2385–2405.

(18) McColgan, P.; Sj, T. Huntington’s disease: a clinical review. Eur J Neurol 2018, 25, 24–34.

(19) Hyeon, C.; Thirumalai, D. Mechanical unfolding of RNA hairpins. Proc. Natl. Acad. Sci. U. S. A. 2005, 102, 6789–6794.

(20) Hyeon, C.; Thirumalai, D. Multiple probes are required to explore and control the rugged energy landscape of RNA hairpins. J. Am. Chem. Soc. 2008, 130, 1538+.

(21) Denesyuk, N. A.; Thirumalai, D. Crowding Promotes the Switch from Hairpin to Pseudoknot Conformation in Human Telomerase RNA. J. Am. Chem. Soc. 2011, 133, 11858–11861.

(22) Denesyuk, N. A.; Thirumalai, D. How do metal ions direct ribozyme folding? Nat. Chem. 2015, 7, 793–801.

(23) Hori, N.; Denesyuk, N. A.; Thirumalai, D. Shape changes and cooperativity in the folding of the central domain of the 16S ribosomal RNA. Proc. Natl. Acad. Sci. U. S. A. 2021, 118.

(24) Hyeon, C.; Dima, R. I.; Thirumalai, D. Pathways and kinetic barriers in mechanical unfolding and refolding of RNA and proteins. Structure 2006, 14, 1633–1645.

(25) Broda, M.; Kierzek, E.; Gdaniec, Z.; Kulinski, T.; Kierzek, R. Thermodynamic stability of RNA structures formed by CNG trinucleotide repeats. Implication for prediction of RNA structure. Biochemistry 2005, 44, 10873–10882.

(26) Sobczak, K.; Michlewski, G.; de Mezer, M.; Kierzek, E.; Krol, J.; Olejniczak, M.; Kierzek, R.; Krzyzosiak, W. J. Structural Diversity of Triplet Repeat RNAs. J. Biol. Chem. 2010, 285, 12755–12764.

(27) Li, Mai Suan and Klimov Dmitri K. and Thirumalai, D., Finite Size Effects on Thermal Denaturation of Globular Proteins. Phys. Rev. Lett. 2004, 93, 268107.

(28) Xu, P.; Pan, F.; Roland, C.; Sagui, C.; Weninger, K. Dynamics of strand slippage in DNA hairpins formed by CAG repeats: roles of sequence parity and trinucleotide interrupts. Nucleic Acids Res. 2020, 48, 2232–2245.

(29) Sobczak, K.; de Mezer, M.; Michlewski, G.; Krol, W. J., Jacek Krzyzosiak RNA structure of trinucleotide repeats associated with human neurological diseases. Nucleic Acids Res. 2003, 31, 5469–5482.

(30) Ni, C.-W.; Wei, Y.-J.; Shen, Y.-I.; Lee, I.-R. Long-Range Hairpin Slippage Reconfiguration Dynamics in Trinucleotide Repeat Sequences. J. Phys. Chem. Lett. 2019, 10, 3985–3990.

(31) Thirumalai, D.; Woodson, S. A. Kinetics of Folding of Proteins and RNA. Accounts of Chemical Research 1996, 29, 433–439.

(32) Treiber, D. K.; Williamson, J. R. Exposing the kinetic traps in RNA folding. Current opinion in structural biology 1999, 9 3, 339–45.

(33) Woodson, S. Recent insights on RNA folding mechanisms from catalytic RNA. Cell. Mol. Life Sci. 2000, 57, 796–808.

(34) Treiber, D.; Williamson, J. Beyond kinetic traps in RNA folding. Curr. Opin. Struct. Biol. 2001, 11, 309–314.

(35) Yoffe, A. M.; Prinsen, P.; Gelbart, W. M.; Ben-Shaul, A. The ends of a large RNA molecule are necessarily close. Nucleic Acids Res. 2010, 39, 292–299.

(36) Xue, Yi and Gracia, Brant and Herschlag, Daniel and Russell, Rick and Al-Hashimi Hashim M, Visualizing the formation of an RNA folding intermediate through a fast highly modular secondary structure switch. Nature communications 2016, 7, comms11768.

(37) Thirumalai, D.; Klimov, D.; Dima, R. Emerging ideas on the molecular basis of protein and peptide aggregation. Curr. Opin. Struct. Biol. 2003, 13, 146–159.

(38) Chakraborty, D.; Straub, J. E.; Thirumalai, D. Differences in the free energies between the excited states of A beta 40 and A beta 42 monomers encode their aggregation propensities. Proc. Natl. Acad. Sci. U. S. A. 2020, 117, 19926–19937.

(39) Ripin, Nina and Parker, Roy, Are stress granules the RNA analogs of misfolded protein aggregates? RNA 2022, 28, 67–75.

(40) Dima, R.; Thirumalai, D. Probing the instabilities in the dynamics of helical fragments from mouse PrPc. Proc. Natl. Acad. Sci. U. S. A. 2004, 101, 15335–15340.

(41) Moulick, R.; Das, R.; Udgaonkar, J. B. Partially Unfolded Forms of the Prion Protein Populated under Misfolding-promoting Conditions: CHARACTERIZATION BY HYDROGEN EXCHANGE MASS SPECTROMETRY AND NMR. J. Biol. Chem. 2015, 290, 25227–25240.

(42) Colon, W.; Kelly, J. Partial denaturation of transthyretin is sufficient for amyloid fibril formation in vitro. Biochemistry 1992, 31, 8654–8660.

(43) Hammarstrom, P.; Wiseman, R.; Powers, E.; Kelly, J. Prevention of transthyretin amyloid disease by changing protein misfolding energetics. Science 2003, 299, 713–716.

(44) c, T.; Cd, S.; Gm, C. Open-to-closed transition in apo maltose-binding protein observed by paramagnetic NMR. Nature 2007, 449, 1078–1082.

(45) Guillaume Bouvignies, G.; Vallurupalli, P.; Hansen, D.; Correia, B.; Lange, O.; Bah, A.; Vernon, R.; W. Dahlquist, F.; Baker, D.; Kay, L. E. Nature 2013, 477, 111–114.

(46) Neudecker, Philipp and Robustelli, Paul and Cavalli, Andrea and Walsh, Patrick and Lundström, Patrik and Zarrine-Afsar, Arash and Sharpe, Simon and Vendruscolo, Michele and Kay, Lewis E, Structure of an intermediate state in protein folding and aggregation. Science 2012, 336, 362—366.

(47) Denesyuk, N. A.; Thirumalai, D. Coarse-Grained Model for Predicting RNA Folding Thermodynamics. J. Phys. Chem. B 2013, 117, 4901–4911.

(48) Denesyuk, N. A.; Hori, N.; Thirumalai, D. Molecular Simulations of Ion Effects on the Thermodynamics of RNA Folding. J. Phys. Chem. B. 2018, 122, 11860–11867.

(49) Oosawa, F. Polyelectrolytes; New York, 1971.

(50) Manning, G. S. Limiting laws and counterion condensation in polyelectrolyte Solutions.I. colligative properties. J. Chem. Phys. 1969, 51, 924–933.

