## Supplemental File for "Odd-even disparity in the population of slipped hairpins in RNA repeat sequences with implications for phase separation"

#### Materials and Methods

##### Model

In the coarse-grained (CG) representation of the RNA sequence, each nucleotide is represented by a single bead, as in the Self-Organized Polymer (SOP) model.[1] The total energy, $E_{TOT}$, is the sum of bonded ($E_B$), hydrogen bonded ($E_{HB}$), excluded volume ($E_{EV}$), and electrostatic energy ($E_{el}$), and is written as,

$$E_{TOT} = E_B + E_{HB} + E_{EV} + E_{el}, \tag{S1}$$

where $E_B$ is the sum of bond stretch ($E_{BS}$) and bond angle ($E_{BA}$) potentials. The expression for $E_B$ is,

$$E_B = E_{BS} + E_{BA} \equiv \frac{1}{2}\sum_{i}^{N_T-1} k_{bond}(r_i - r_0)^2 + \frac{1}{2}\sum_{j}^{N_T-2} k_{angle}(\alpha_j - \alpha_0)^2, \quad \text{(S2)}$$

where $N_T$ is the number of nucleotides in a given sequence. The parameters, $k_{bond}$ and $k_{angle}$, are the spring constants in $E_{BS}$ and $E_{BA}$, respectively. In Eq. S2, $r_i$ and $\alpha_j$ are the $i^{th}$ bond distance and $j^{th}$ bond angle, respectively, and $r_0$ and $\alpha_0$ are the corresponding equilibrium values.

Hydrogen bond interactions between two beads, with complementary base pairs such as cytosine (C) and guanine (G), and, adenine (A) and uracil (U), contribute substantially to the stability of the folded hairpin-like structures that the low complexity sequences adopt. The functional form for $E_{HB}$ is,

$$E_{HB} = N_b u_{hb}^0 \exp[-U_{hb,bond} - U_{hb,angle} - U_{hb,dihedral}], \quad \text{(S3)}$$

where $N_b = 3$ (2) for a G-C (A-U) base pair (bp), and $u_{hb}^0$ is the strength of a single hydrogen bond. We varied $u_{hb}^0$ and calculated the heat capacities ($C_v(T)$s) for a few $n$ values. The peak in $C_v(T)$ is associated with the melting temperature, $T_M$. The value of $u_{hb}^0$ is determined by matching the experimental and calculated melting temperatures of $G(CAG)_nC$ sequences for $n = 5$, 6 and 7 as closely as possible. The temperature dependent $C_v(T)$s in Fig.S1 show that $u_{hb}^0 =$ -2.0 kcal/mol gives good agreement between simulations and experiments for melting temperatures for $n$=6 and 7. Without further refinement, we performed all simulations with this value of $u_{hb}^0$.

The expressions for $U_{hb,bond}$, $U_{hb,angle}$ and $U_{hb,dihedral}$ are,

$$U_{hb,bond} = k_r(r_{ij} - r_{hb,0})^2, \quad \text{(S4)}$$

$$U_{hb,angle} = k_\theta(\theta_{i,j,j-1} - \theta_1)^2 + k_\theta(\theta_{i-1,i,j} - \theta_1)^2 + k_\theta(\theta_{i,j,j+1} - \theta_2)^2 + k_\theta(\theta_{i+1,i,j} - \theta_2)^2, \quad \text{(S5)}$$

$$U_{hb,dihedral} = k_\phi[1 + cos(\phi_{j-1,j,i,i-1} + \phi_1)] + k_\phi[1 + cos(\phi_{j+1,j,i,i+1} + \phi_2)]. \quad \text{(S6)}$$

In Eq. S4, $r_{ij}$ is the distance between beads $i$ and $j$, $r_{hb,0}$ is the hydrogen bond distance in an ideal A-form helix, and $k_r$ is a constant. In Eq. S5, $\theta_{i,j,k}$ is the angle between beads $i$, $j$ and $k$, and $\theta_1$ and $\theta_2$ are two constants. Contribution of the angular part to the total hydrogen bond potential is determined by $k_\theta$, which has the unit of radian$^{-2}$. In Eq. S6, the dihedral angle, $\phi_{i,j,k,l}$, is the angle between the two planes defined by the four beads $i$, $j$, $k$ and $l$, and, $k_\phi$, $\phi_1$ and $\phi_2$ are constants.

In order to avoid unphysical overlap between two non-bonded beads that are separated by two other beads, we used a repulsive potential,

$$E_{EV} = \epsilon_{ev}[(\frac{\sigma}{r_{ij}})^{12} - (\frac{\sigma}{r_{ij}})^6 + 1] \text{ if } r_{ij} \le \sigma, \quad \text{(S7)}$$

$$E_{EV} = 0 \text{ otherwise}, \quad \text{(S8)}$$

where, $\epsilon_{ev}$ is the strength of repulsive interaction, and $\sigma = 10$ Å. The values of the parameters in the energy function are listed in Table S1.

#### Electrostatic interactions and charge renormalization

In the presence of cations, the charge on each bead is reduced to -Q ($0 < Q < 1$) from -1 due to counterion condensation.[2,3] The effects of monovalent salt ions are included implicitly in the SOP model by using a renormalized $Q$ value. We calculated $Q$, in presence of monovalent salts, as $\frac{b}{l_B(T)}$, where $b$ (=4.4 Å)[4] is the length per unit charge. The Bjerrum length, $l_B(T) = \frac{e^2}{\epsilon(T)k_BT}$ where $e$ is the electron charge, $\epsilon(T)$ is the temperature dependent dielectric constant of water, $k_B$ is the Boltzmann constant, and $T$ is the temperature. The temperature dependent $\epsilon(T)$, $T$ (in °C) is computed using,[5]

$$\epsilon(T) = 87.74 - 0.4008T + 9.398 * 10^{-4}T^2 - 1.410 * 10^{-6}T^3. \quad \text{(S9)}$$

The electrostatic energy between two negatively charged phosphate groups is taken to be,

$$E_{el} = \frac{Q^2 e^2}{\epsilon(T)} \sum_{i<j} \frac{\exp(-\kappa_D r_{ij})}{r_{ij}}, \quad \text{(S10)}$$

where $\kappa_D$ $(=\sqrt{8\pi\rho l_B(T)})$ is the inverse Debye screening length. In monovalent salt solutions $q_i = 1$ is the magnitude of the charge on each counterion and $\rho$ is the number density of ions in the solution. For a given monovalent salt concentration, $C_s$, it is easy to show that $\rho = 2C_s \times 6.022 \times 10^{-4}$ Å$^{-3}$.

#### Inter-chain interactions in dimers:

Interactions between nucleotides belonging to two RNA chains are the same as in intra-chain interactions. In order to maintain proximity between the two chains so that dimers form on reasonable time scales in the simulations, the center of mass between the chains are constrained by a harmonic potential, $U_D$,[6]

$$U_D = \frac{1}{2} k_D (R_{cm} - R_0)^2 \text{ if } R_{cm} > R_0. \quad \text{(S11)}$$

If $R_{cm} < R_0$ then we set $U_D = 0$ . In Eq.S11, $R_{cm}$ is the center of mass distance between the two chains, $R_0$ is the cutoff distance. We chose $R_0 \approx \langle R_g \rangle$ of the monomers to mimic the concentration for the dimer formation. The value of $R_0 = 40$ Å when dimer formation is initiated from the ground state. We used $R_0 = 100$ Å if the initial structures correspond to unfolded polynucleotides. The value of $k_D$ is 200 kcal/(mol.Å$^2$). Note that for the restraining potential in Eq. S11 the precise value of $R_0$ has negligible effect on dimer formation rates or yields.

#### Simulations:

To extract the thermodynamics of $(CAG)_n$, we performed low friction Langevin dynamics simulations[7] by integrating the equation motion, which for bead $i$ is,

$$m\ddot{\vec{r_i}} = -\zeta\dot{\vec{r_i}} + \vec{F_C} + \vec{\Gamma}. \tag{S12}$$

In the above equation, $m$ is the mass of each bead, $\zeta$ is the friction coefficient, $F_C = -\frac{\partial E_{TOT}}{\partial \vec{r_i}}$, and $\vec{\Gamma}$ is the random force with Gaussian statistics, $\langle \Gamma(t)\, \Gamma(t+ph) \rangle = \frac{2\zeta k_B T}{h}\delta_{0,p}$ where $p = 0, 1, \cdots$, $\delta_{0,p}$ is the Kronecker delta function. We integrated Eq. S12 using the velocity Verlet algorithm. Following our previous work,[7] we used a small value of the friction coefficient in order to sample the conformational space efficiently. The friction coefficient $\zeta = 0.05\ \frac{m}{\tau_L}$ and $h = 0.005\ \tau_L$, where $\tau_L = (\frac{m_0 a_0^2}{\epsilon_h})^{0.5}$ is the time unit in the Langevin dynamics simulations. Thermodynamic properties do not depend on the precise value of $\zeta$ or $m$.

To investigate dimer formation, we performed Brownian dynamics simulations using the Ermak-McCammon algorithm,[8]

$$\dot{\vec{r_i}}(t+h) = \vec{r_i}(t) + \frac{h}{\zeta}\vec{F_C}(t) + \vec{R}(t), \tag{S13}$$

where $\vec{R}(t)$ is the random displacement with mean $\langle R(h) \rangle = 0$, and variance $\langle [R(h)]^2 \rangle = \frac{2k_B T h}{\zeta}$. The time step, $h$, to integrate the equation of motion is taken as $0.05\tau_H$, where, $\tau_H$ is the time unit of Brownian dynamics simulations and $\zeta$ is taken as 2.6 $m/\tau_H$. The value of $\zeta$ used in our simulations is $\approx 10$ times smaller compared to the actual friction coefficient of the water. However, we use this lower value in order to observe dimer formation in a reasonable time. The value of $\tau_H = \frac{\zeta_H a_0^2}{k_B T} \approx 40$ ps. Thus, the rates obtained from the simulations are not quantitative, and are lower bounds to the actual rates.

#### Weighted Histogram Analysis Method (WHAM)

All the thermodynamic properties ($\langle A(T) \rangle$ were calculated using the Weighted Histogram Analysis Method,[9]

$$\langle A(T) \rangle = Z(T)^{-1} \sum_{k=1}^{R} \sum_{t=1}^{n_k} \frac{A_{k,t} \exp(-E_{k,t}/k_B T)}{\sum_{m=1}^{R} n_m \exp(f_m - E_{k,t}/k_B T m)}, \tag{S14}$$

where $R$ is the number of simulation trajectories, $n_k$ is the total number of conformations in the $k^{th}$ simulation, $A_{k,t}$ is the value of $A$ for $t^{th}$ conformation in the $k^{th}$ simulated trajectory, $E_{k,t}$ is the potential energy for conformation $t$ in trajectory $k$, $T_m$ and $f_m$ are the temperature and free energy of trajectory $m$, respectively. The partition function, $Z(T)$, is,

$$Z(T) = \sum_{k=1}^{R} \sum_{t=1}^{n_k} \frac{\exp(-E_{k,t}/k_B T)}{\sum_{m=1}^{R} n_m \exp(f_m - E_{k,t}/k_B T_m)}. \tag{S15}$$

#### Order parameter, $\chi$:

Structural similarity of a conformation, with respect to a reference structure, is quantified using the overlap parameter, $\chi$,[10] which is defined as,

$$\chi = \frac{1}{N_P} \sum_{i,j}^{N_P} \Theta(d - |r_{ij} - r_{ij}^0|), \tag{S16}$$

where $N_P$ is the number of non-bonded pairs of beads, $r_{ij}$ is the distance between bead $i$ and $j$ and $r_{ij}^0$ is the corresponding distance in the reference structure taken from the Protein Data Bank (PDB) entry, and $\Theta$ is the Heaviside step function, and $d$, the tolerance = 2.5 Å.

### Folding Thermodynamics

#### Unfolded to folded transitions occur in a two-state manner

In order to establish the nature of the thermodynamic transition, we calculated the distribution of the structural overlap function $\chi$ (Equation S16), which requires knowledge of the ground state of the RNA sequences. Rather than use the simulated ground state, we used the crystal structures for $(CAG)_2$. The coordinates for $G(CAG)_6C \equiv G(CAG)_2(CAG)_2(CAG)_2C$ are obtained by stitching together the three $(CAG)_2$ units. For each $(CAG)_2$ we used the available crystal structure (protein data bank (PDB) (entry 3NJ7.pdb)). Except for the terminal nucleotide at the $5'$ and $3'$ ends, the primary sequence of the duplex is the same as the stem region in the hairpin formed by the $G(CAG)_6C$ sequence (Figure S3). Using the PDB structure, we produced a reference structure for the GS for $G(CAG)_6C$ needed to calculate $\chi$. The temperature dependence of the stem region of the $G(CAG)_6C$ sequence exhibits a sigmoidal shape (Figure S4A), indicating that $G(CAG)_6C$ folds in an apparent two state manner. Although the probability distribution $P(\chi)$ is broad, (Figure S4B) at $T_M$ it is approximately bimodal, which supports the approximation that thermal folding occurs by a two-state transition.

#### Fraction of base pairs versus $T$

To probe the structural changes as the temperature is increased, we calculated the average fraction of base pairs, $\langle f_{bp} \rangle$ ($f_{bp} = \frac{N_{bp}}{N_{bp}^{max}}$, ($N_{bp}$ is the number of base pairs in a conformation, and $N_{bp}^{max}$ is the number of base pairs in a perfect hairpin conformation), as a function of temperature at $C_s = 1.0$ M for $G(CAG)_nC$ with $n = 5$, 6 and 7 and $C_s = 0.1$ M for $(CAG)_n$ with $n = 14$, 15 and 20 (Figure S5). Here after, $C_s = 1.0$ M for $G(CAG)_nC$ with n = 5, 6 and 7 and $C_s = 0.1$ M for $(CAG)_n$ with n= 14, 15, 20, 30 and 31 is denoted as standard salt concentration. For all the sequences, $\langle f_{bp} \rangle$ changes from $\approx 0.8$ to $\approx 0$ as the temperature increases from 27°C to 127°C (Fig. S5).

To classify the conformations either as folded or unfolded, we used a critical value for the fraction of base pairs, $f^c_{bp}$=0.5. Conformations with $f_{bp} \geq f^c_{bp}$ are classified as folded state. Otherwise, they are assumed to be unfolded.

#### Free energy difference, $\Delta G = G_f - G_u$

We determined $\Delta G$ by calculating the temperature dependence of the free energies of the folded ($G_f$) and unfolded states ($G_u$) using a procedure described previously.[4] The free energy of the folded state, $G_f(T)$, is calculated using,

$$G_f(T) = G(T^*_f) + \frac{\partial G}{\partial T}(T^*_f)(T - T^*_f) + T\frac{\partial^2 G}{\partial T^2}(T^*)\left(T^*_f - T + T\ln\frac{T}{T^*_f}\right). \tag{S17}$$

We chose a sufficiently low value of $T^*$ such that the RNA is stably folded. Similarly, the $G_u$ is computed by choosing a high temperature threshold, $T^*_u$, above which the the unfolded state is stable. In our simulations, $T^*_f$ is in the range 0° C to 30° C and $T^*_u$ is varied between 100° C to 130° C depending on the sequence length. We calculated the free energy $G_\alpha(T)$ ($\alpha$ = u or f) using, $G_\alpha(T) = -k_B T \ln Z_\alpha(T)$, where $k_B$ is Boltzmann constant and $Z_\alpha(T)$ is the partition function of the $\alpha^{th}$ state (Eq. S15). The temperature dependencies of $\Delta G(T)$ for various sequences are shown in Fig.S6. The melting temperature could be estimated using $\Delta G(T_M) = 0$. Values of $T_M$ may be obtained using the relation $\langle f_{bp}\rangle = 0.5$ (Fig. S5). The two ways of computing $T_M$ yield similar values.

#### Salt effects on the thermodynamics of $(CAG)_n$ repeats

To quantify salt effects on the stability of the folded state, we calculated the changes in free energy difference, $\Delta\Delta G$ using $\Delta\Delta G = \Delta G(C_s) - \Delta G(C_0)$, where $\Delta G(C_s) = G_f$ - $G_u$ at $C_s$ and $\Delta G(C_0)$ is evaluated at $C_0$ (Fig. S7). We use $C_0$ = 0.15 M as the reference salt concentration because it roughly corresponds to the physiological value. We calculated $\Delta\Delta G$ at $T$=27°C and 37°C for all the sequences except for $AG(CAG)_5CA$ for which $T$ was set

2.5°C and 27°C. Fig. S7 shows that $\Delta\Delta G$ decreases linearly with $C_s$ with a negative slope at $C_s < 0.2$ M and deviates from linearity at high $C_s$.[4]

#### Structures with fractional $Q_{HP}$

In the main text we showed schematically (Figure 2) conformations with integer values of the order parameter $Q_{HP}$ (defined in Eq. (1) in the main text). Examples of structures with non-integer values of $Q_{HP}$ for A(CAG)$_{14}$A and A(CAG)$_{15}$A are shown in Fig. S8. The structures with fractional values of $Q_{HP}$ typically have bulges with unsatisfied G-C base pairs. As a consequence, they must correspond to excited states of the low complexity sequences (see below).

#### Free energy spectra

The free energy spectra, relative to the ground states, for even and odd $n$, calculated using Eq. (2) in the main text are displayed in Fig. S9 and Fig.S10, respectively. The distributions $P(Q_{HP}$s are shown in Figure 2A ($n$ even) and Figure 2B ($n$ odd) in the main text. Interestingly, the relative free energy of the state with $Q_{HP} = 2$ is lower than $Q_{HP} = 1$ when $n$ is even (Fig. S9). An explanation is found in their conformations. In the loop region of A(CAG)$_{14}$A there are four nucleotides when $Q_{HP} = 2$ whereas seven nucleotides (heptaloop) are required to generate conformations with $Q_{HP} = 1$ (see Figure 2C in the main text). Loss in loop entropy makes the free energy of the $Q_{HP} = 1$ state marginally higher (Fig. S9). Although the free energy loss due to slippage (2 CAG units in $Q_{HP} = 2$) is greater than in conformations with $Q_{HP} = 1$ (one CAG slip), it is apparently compensated by the formation of a shorter loop. Of course, the difference between the free energies between the $Q_{HP} = 1$ and $Q_{HP} = 2$ states is marginal (less than $k_BT$). When $n$ is odd, the PHs are the excited states (FigureS10 with free energies that are greater than $k_BT$ compared to the ground state with one slipped CAG (Fig. 2D in the main text).

### Dimerization from unfolded state

#### Dimerization pathways depend on initial conditions

We initiated dimer simulations starting from the unfolded conformations for $(CAG)_{30}$ and $(CAG)_{31}$ at $C_s = 0.1$ M and $T = 37°$. Dimers form in $\approx 30$ % of the trajectories. The time dependent changes in the loss of intramolecular base pairs and gain in the intermolecular base pairs are shown in Fig. S13A and Fig. S13B. In majority of such trajectories dimerization occurs directly from the unfolded configurations (Fig. S13A). In a small fraction of the trajectories, the formation of hairpin in the two chains precedes dimerization (Fig. S13C). A schematic representation of dimer formation pathways starting from the unfolded state is shown in Fig. S13C. Interestingly, there is essentially no difference in the rate of dimer formation between $A(CAG)_{30}A$ and $A(CAG)_{31}A$ (Fig. S13D), which shows that the mechanism of self-association between RNA chains depends on the starting conditions.

#### Exact sequence matters for self-association

Recent experiments[11] have probed the effects of sequences in the self-association of RNA polymers. Although multivalent interactions are important, the association mechanism of chains with identical number of multivalent sites but variation in sequences could be different.[12] To investigate the role of sequence in determining dimer formation, we compared the rates of dimerization for two mutants of M3 (sequence, $A(CAG)(CAG)_{28}(CCG)_3A$) and M4 (sequence, $A(CAG)_{12}(CCG)_3CAG(CGG)_3(CAG)_{12}A$), which have the same composition and length. The M3 sequence in which the CC and GG rich region is located near the termini forms dimer at $\approx 2.2$ times faster compared to M4 in which the CC and GG rich region is around the middle of the chain (Fig. S14B). Proximity of the CC and GG rich regions in M4 enhances intra-molecular base pair formation, thus decreasing the rate of dimerization. Our results show that, besides multivalency, the excitation spectra, which is a finger print of the sequence and environmental conditions, is vital in governing the aggregation tendency of RNA chains.

Table S1: List of parameters used in the single bead SOP energy function

| | |
|---|---|
| $k_{bond}$ | 15 kcal/mol. Å$^{-2}$ |
| $r_0$ | 5.9 Å |
| $k_{angle}$ | 10 kcal/mol.rad$^{-2}$ |
| $\alpha_0$ | 2.618 rad |
| $\epsilon_{EV}$ | 2.0 kcal/mol |
| $\sigma$ | 10 Å |
| $r_{hb,0}$ | 13.8 Å |
| $k_r$ | 3.0 Å$^{-2}$ |
| $k_\theta$ | 1.5 rad $^{-2}$ |
| $k_\phi$ | 0.5 |
| $\theta_1$ | 1.8326 rad |
| $\theta_2$ | 0.9425 rad |
| $\phi_1$ | 1.8326 rad |
| $\phi_2$ | 1.345 rad |
| $u_{hb,0}$ | -2.0 kcal/mol |

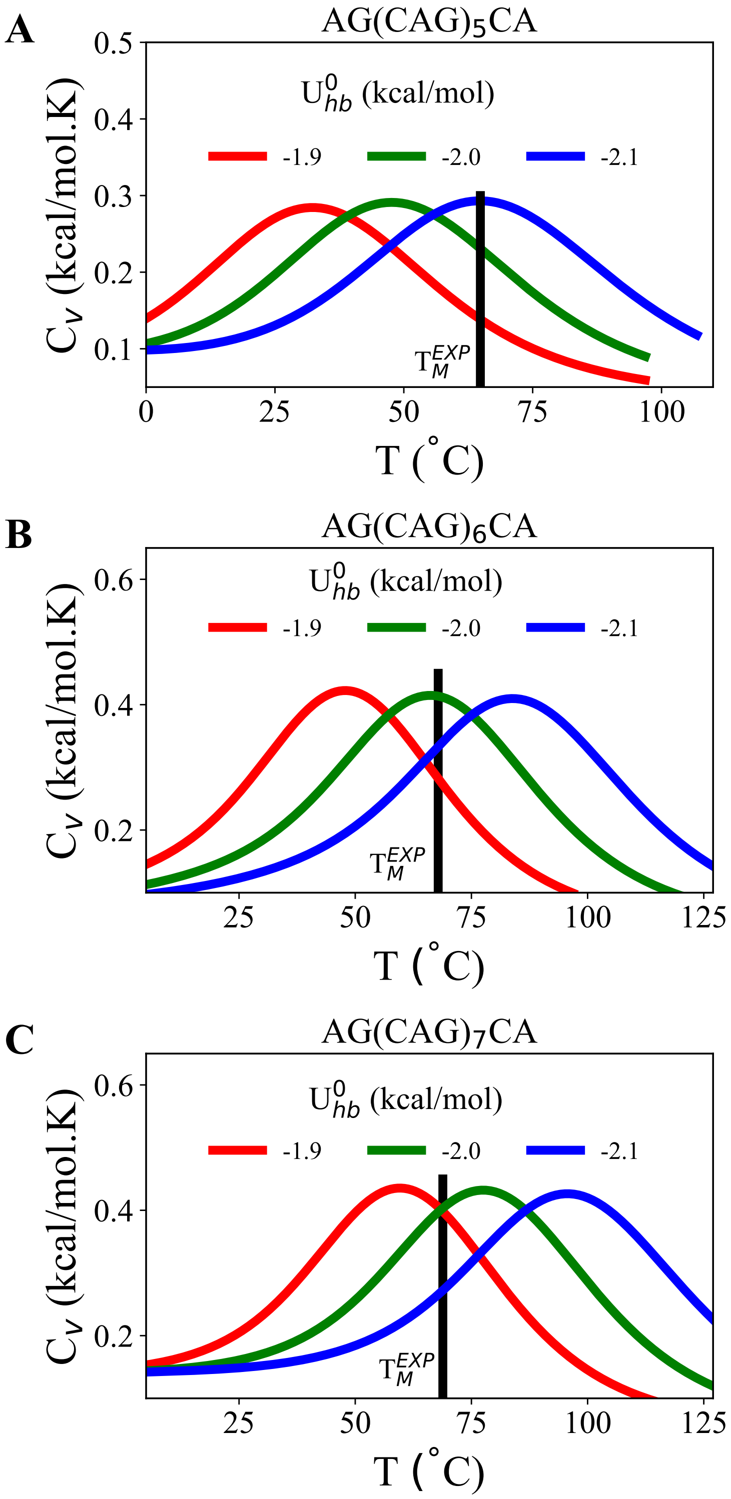


Figure S1: **Choosing $u^0_{hb}$.** Temperature-dependent heat capacities ($C_v(T)$) for (A) G(CAG)$_5$C, (B) G(CAG)$_6$C, and (C) G(CAG)$_7$C with $u^0_{hb}$ = -1.9 (red), -2.0 (green) and -2.1 kcal/mol (blue), respectively. Experimental melting temperatures, $T_M$, obtained at $C_s$ = 1 M are shown by the vertical lines.

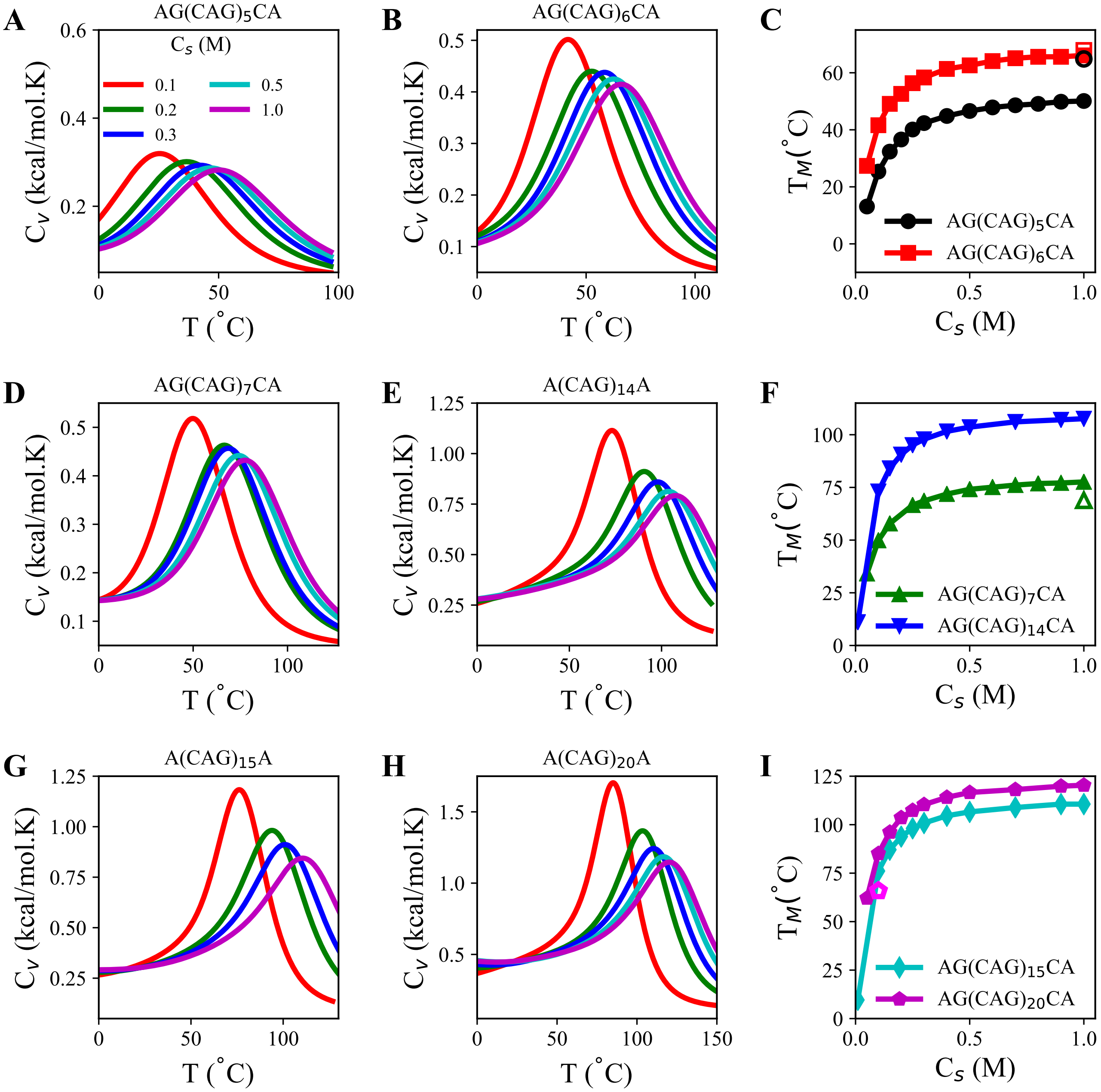


Figure S2: **Thermodynamics of melting.** Heat capacity $C_v$ as a function of temperature at $C_s$ = 0.1 M (red), 0.2 M (green), 0.3 M (blue), 0.5 M (cyan) and 1.0 M (magenta) (A) G(CAG)$_5$C, (B) G(CAG)$_6$C, (D) G(CAG)$_7$C, (E) (CAG)$_{14}$, (G) (CAG)$_{15}$, (H) (CAG)$_{20}$. (I) Melting temperature, $T_M$ as a function of $C_s$: (C) G(CAG)$_5$C and G(CAG)$_6$C are in black circles and red squares, respectively. Experimental melting temperature, $T_M^{EXP}$, at $C_s$ = 1 M for G(CAG)$_5$C and G(CAG)$_6$C are in open black circle and open red square, respectively. (F) G(CAG)$_7$C and (CAG)$_{14}$ are in green triangles and blue inverted triangles, respectively. $T_M^{EXP}$ for G(CAG)$_7$C is in open green triangle. (I) (CAG)$_{15}$ and (CAG)$_{20}$ are in cyan diamonds and magenta pentagons, respectively. $T_M^{EXP}$ for (CAG)$_{20}$ is in open magenta pentagon.

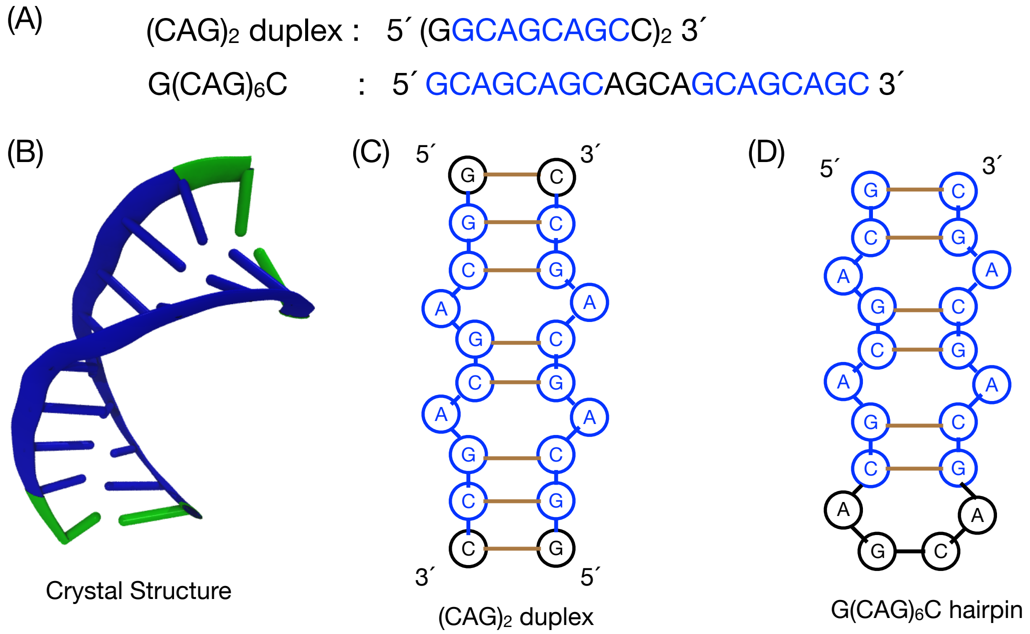


Figure S3: **Sequences and structures for $(CAG)_2$ and $(CAG)_6$.** (A) Primary sequence of $(CAG)_2$ and $G(CAG)_6C$. Sequence in blue is used to calculate structural overlap parameter, $\chi$ (Eq. 5 in the main text). (B) Crystal structure of $(CAG)_2$ duplex (Protein Data Bank (PDB) id 3nj7.pdb) rendered with visual molecular dynamics (VMD). (C) Schematic representation of the secondary structure of $(CAG)_2$ duplex. (D) Same as (C) except this is for $G(CAG)_6C$ hairpin. Sequence in black forms the loop and the the ones in blue are the stems.

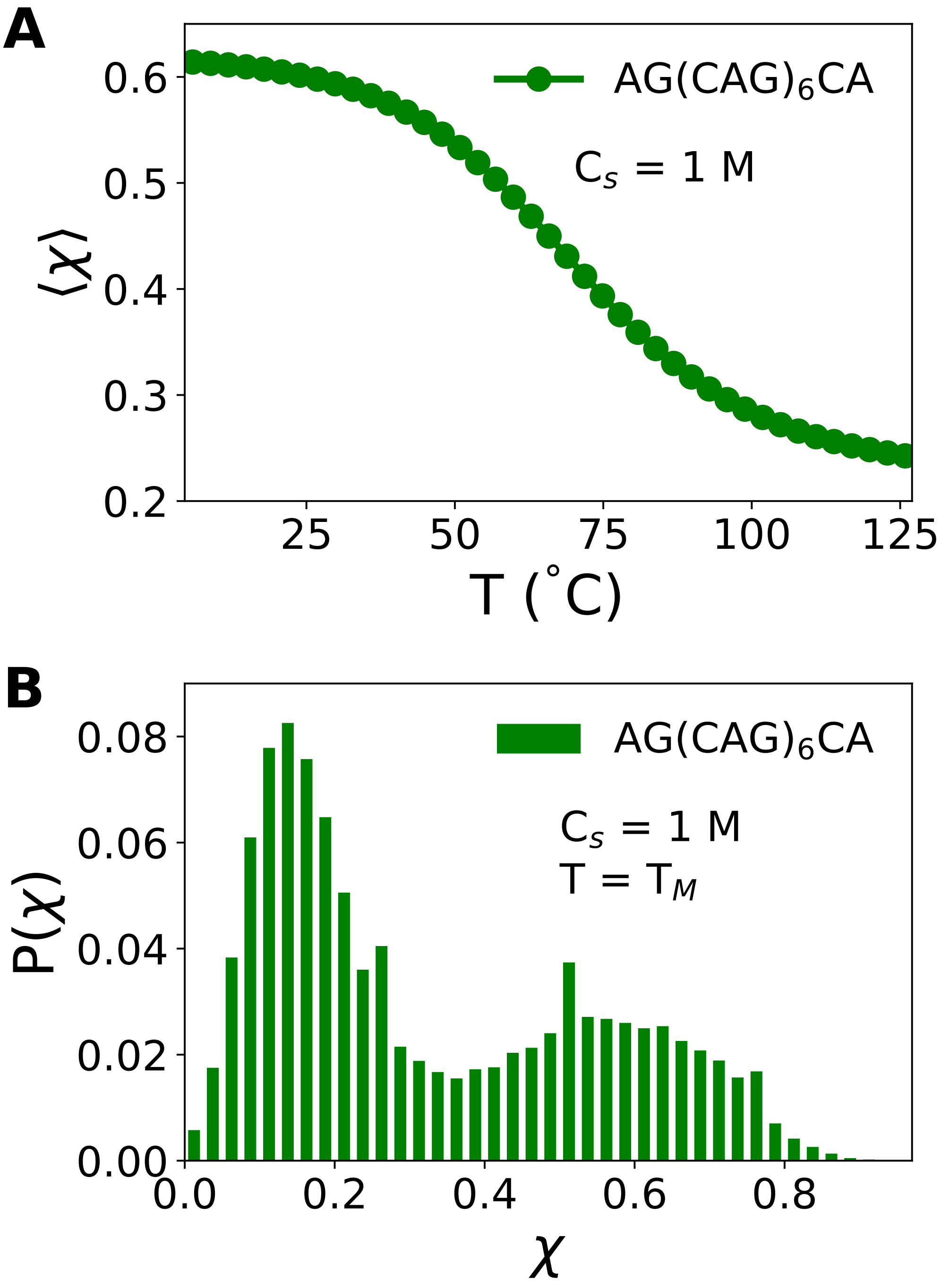


Figure S4: **Approximate two-state transition.** (A) Average of the structural overlap parameter, $\langle\chi\rangle$ (Eq. S16), as a function of temperature for the stem region of G(CAG)$_6$C hairpin. (B) Probability distribution (P($\chi$)) as a function of $\chi$, (P($\chi$)), for G(CAG)$_6$C at the melting temperature, $T_M$ and salt concentration, $C_s = 1$ M. Despite the small stem size, P($\chi$) exhibits bimodal distribution with a minimum at $\chi \approx 0.37$ that separates the folded and unfolded states.

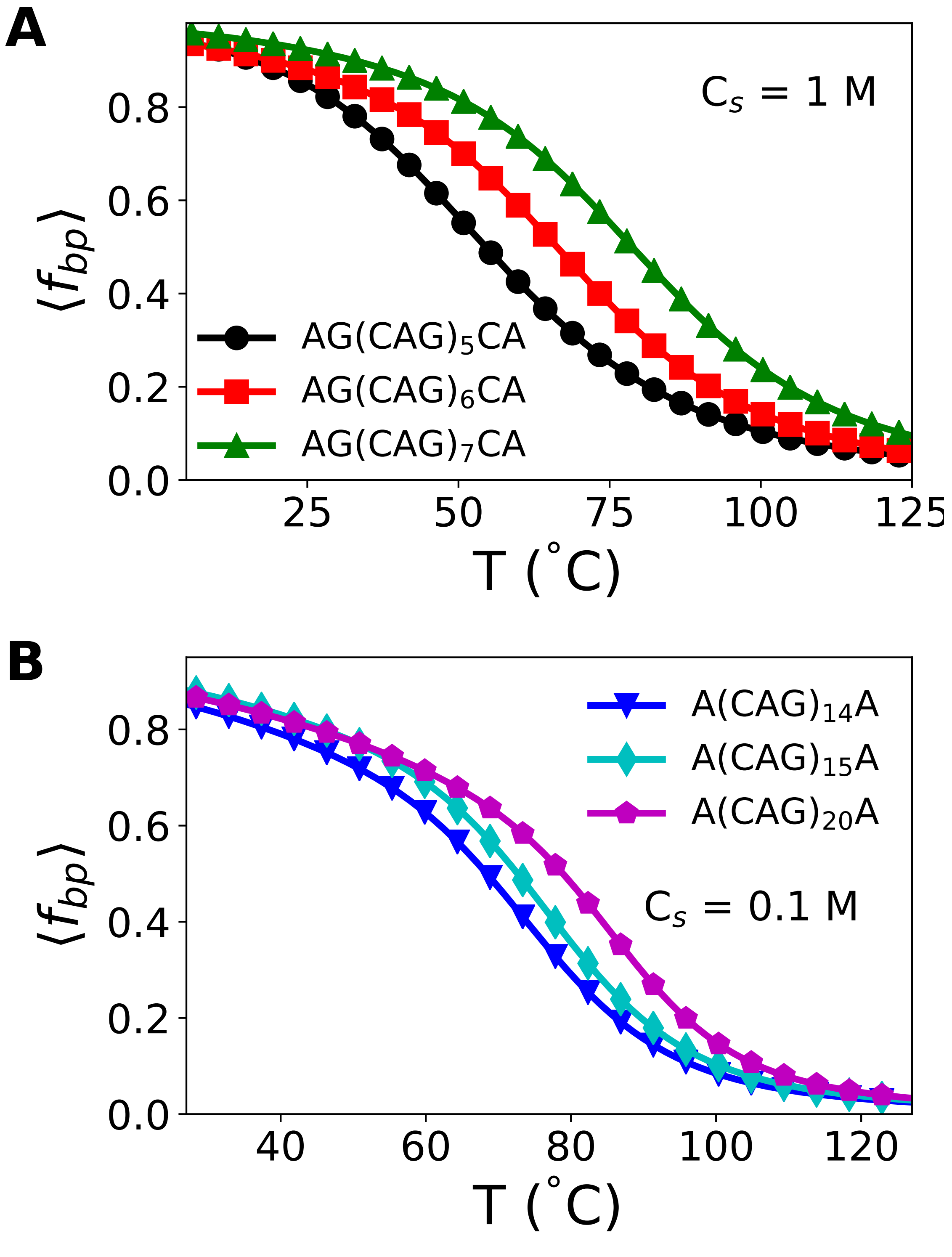


Figure S5: **Average fraction of base pairs, $\langle f_{bp} \rangle$, as a function of temperature.** (A) G(CAG)$_5$C, G(CAG)$_6$C and G(CAG)$_7$C are in black circles, red squares and green triangles, respectively. (B) $\langle f_{bp} \rangle$ for (CAG)$_{14}$, (CAG)$_{15}$ and (CAG)$_{20}$ are in blue inverted triangles, cyan diamonds and magenta pentagons, respectively. Stability of the folded state increases as $n$ increases.

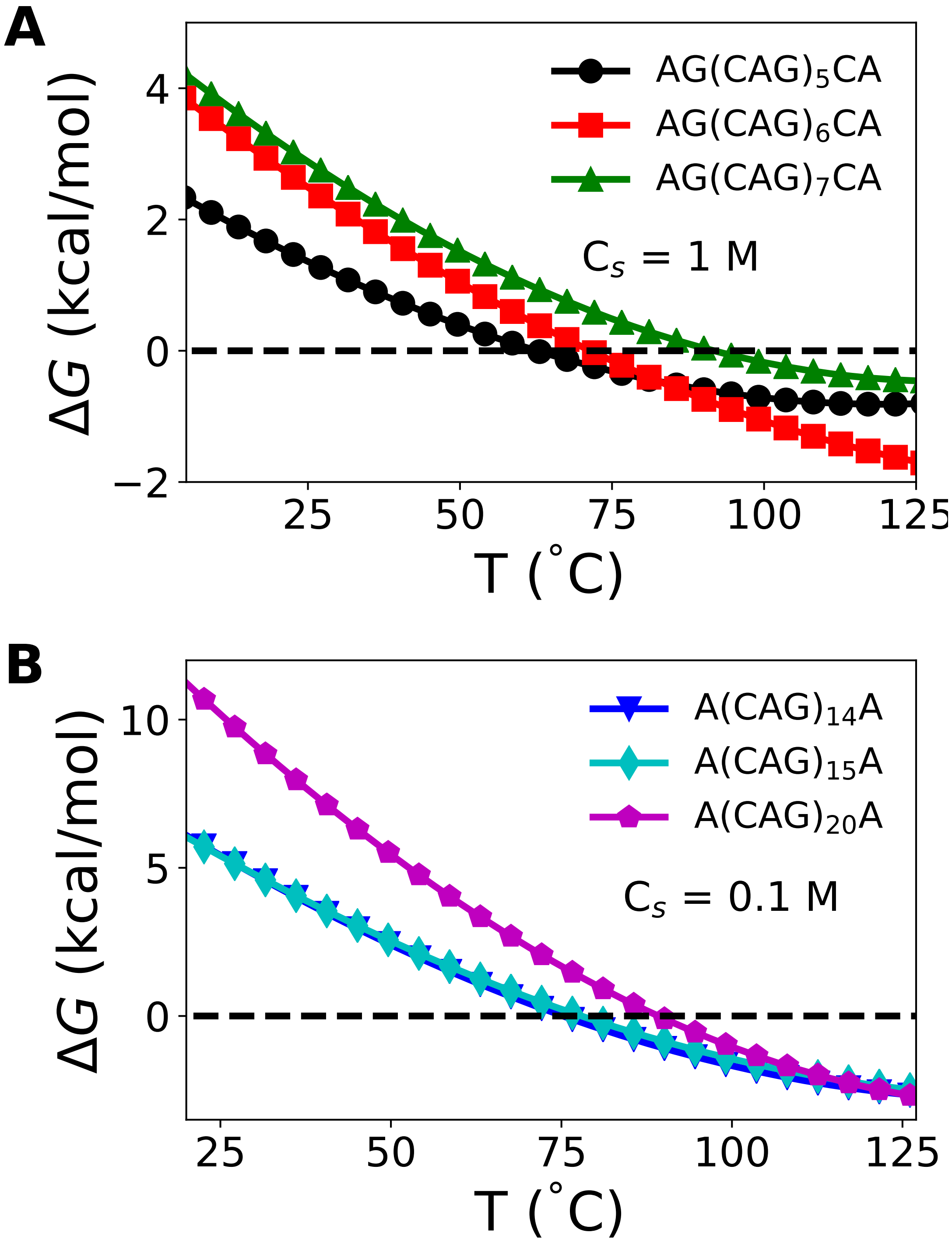


Figure S6: **Sequence-dependent stabilities as a function of $T$:** Free energy difference, $\Delta G = G_u - G_f$, where $G_u$ ($G_f$) is the free energies of the unfolded (folded) state as a function of temperature: (A) G(CAG)$_5$C, G(CAG)$_6$C, G(CAG)$_7$C are in black circles, red squares and green triangles, respectively. (B) (CAG)$_{14}$, (CAG)$_{15}$ and (CAG)$_{20}$ are in inverted blue triangles, cyan squares and magenta pentagons, respectively. Black dashed line ($\Delta G = 0$) separates the folded and the unfolded states.

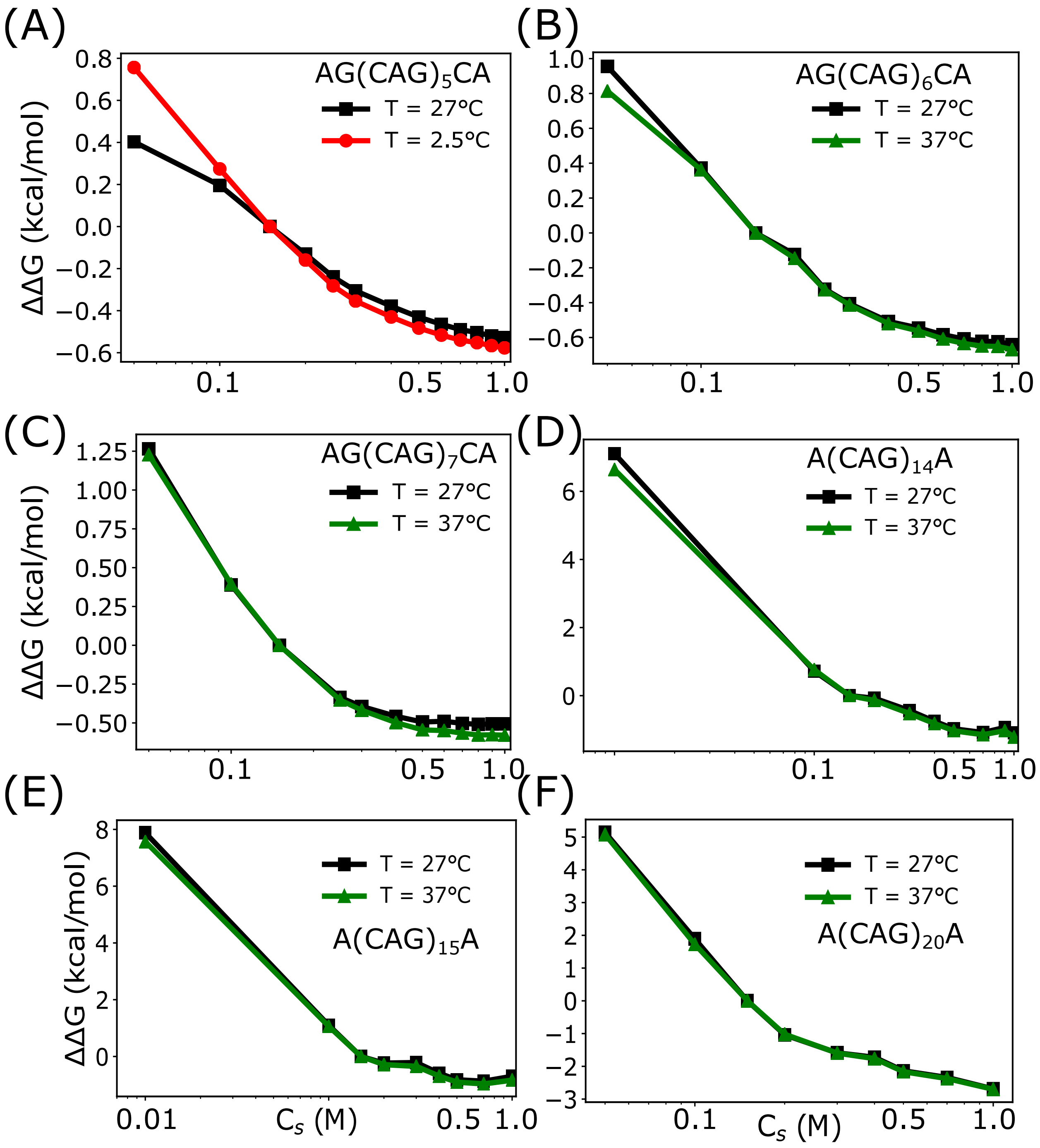


Figure S7: **Free energy difference, $\Delta\Delta G = \Delta G(C_s) - \Delta G(0.15M)$ as a function of $C_s$.** (A) G(CAG)$_5$C , (B) G(CAG)$_6$C, (C) G(CAG)$_7$C, (D) (CAG)$_{14}$, (E) (CAG)$_{15}$, and (F) (CAG)$_{20}$ at different temperatures. $\Delta\Delta G$ values at 2.5°C, 27°C and 37°C are in red circles, black squares and green triangles, respectively.

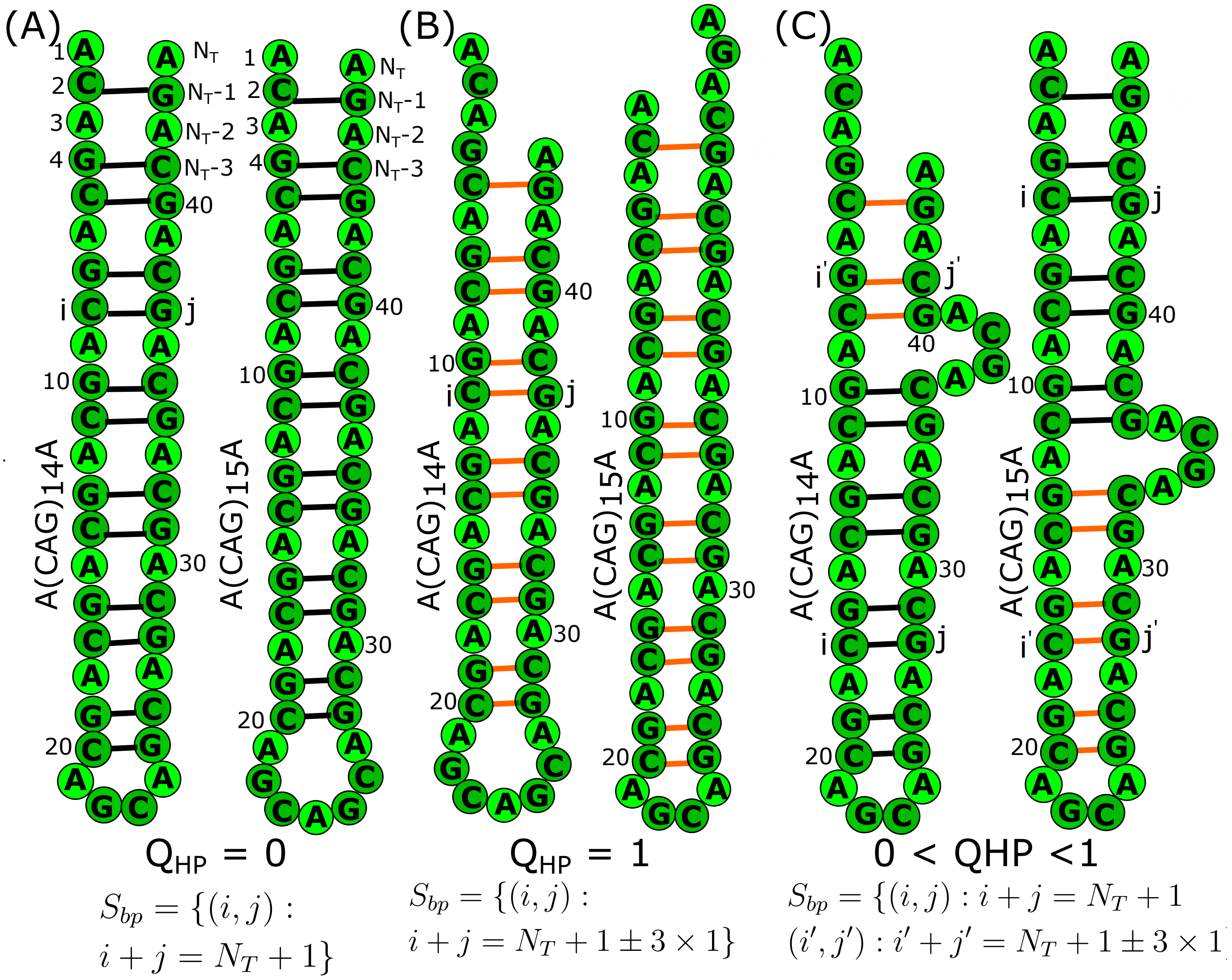


Figure S8: **Secondary structures maps for sequences with even and odd number of repeats.** (A) Hairpin structures for $(CAG)_{14}$ and $(CAG)_{15}$ in which the strands are perfectly aligned, resulting in a perfect hairpin with $Q_{HP} = 0$. To obtain $Q_{HP} = 0$ for $(CAG)_{15}$ the number of nucleotides in the loop has to be 7. (B) Hairpins in $(CAG)_{14}$ and $(CAG)_{15}$, with deviation from a PH due to slippage in strands at $3'$ and $5'$ end, respectively. The value of $Q_{HP}$ is unity. (C) Hairpin structures with a fractional value of $Q_{HP}$ containing. Subset of base pairs in black is identical to $Q_{HP} = 0$, whereas another subset in orange is identical to $Q_{HP} = 1$. In all structures, $S_{bp}$ is expressed in terms of the nucleotide indices $i$ and $j$ that form base pairs.

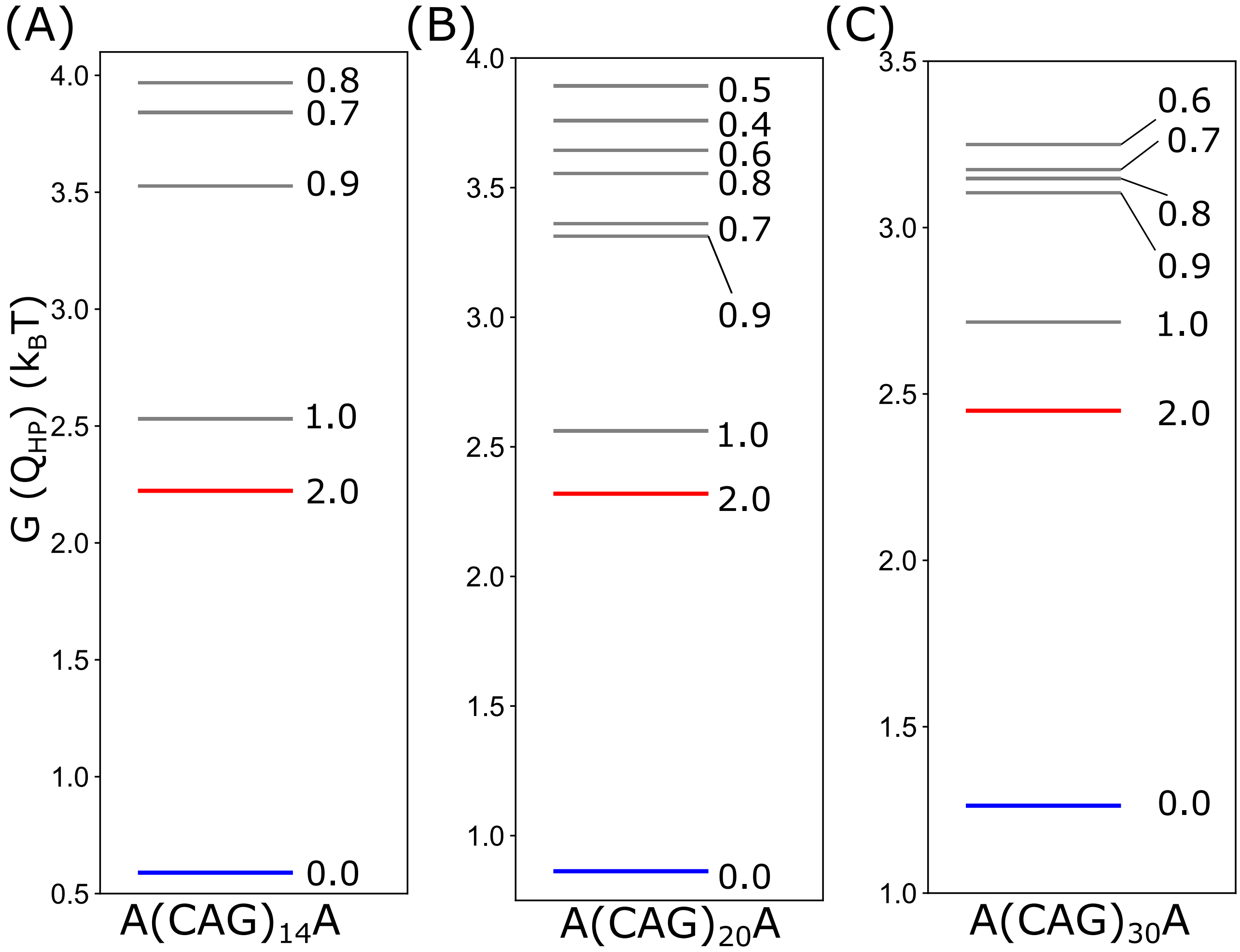


Figure S9: **Free energy spectrum, $G(Q_{HP})$ for even $n$ as a function of $Q_{HP}$.** (A) $G(Q_{HP})$ for A(CAG)$_{14}$A. (B) $G(Q_{HP})$ for A(CAG)$_{20}$A. (C) $G(Q_{HP})$ for A(CAG)$_{30}$A.

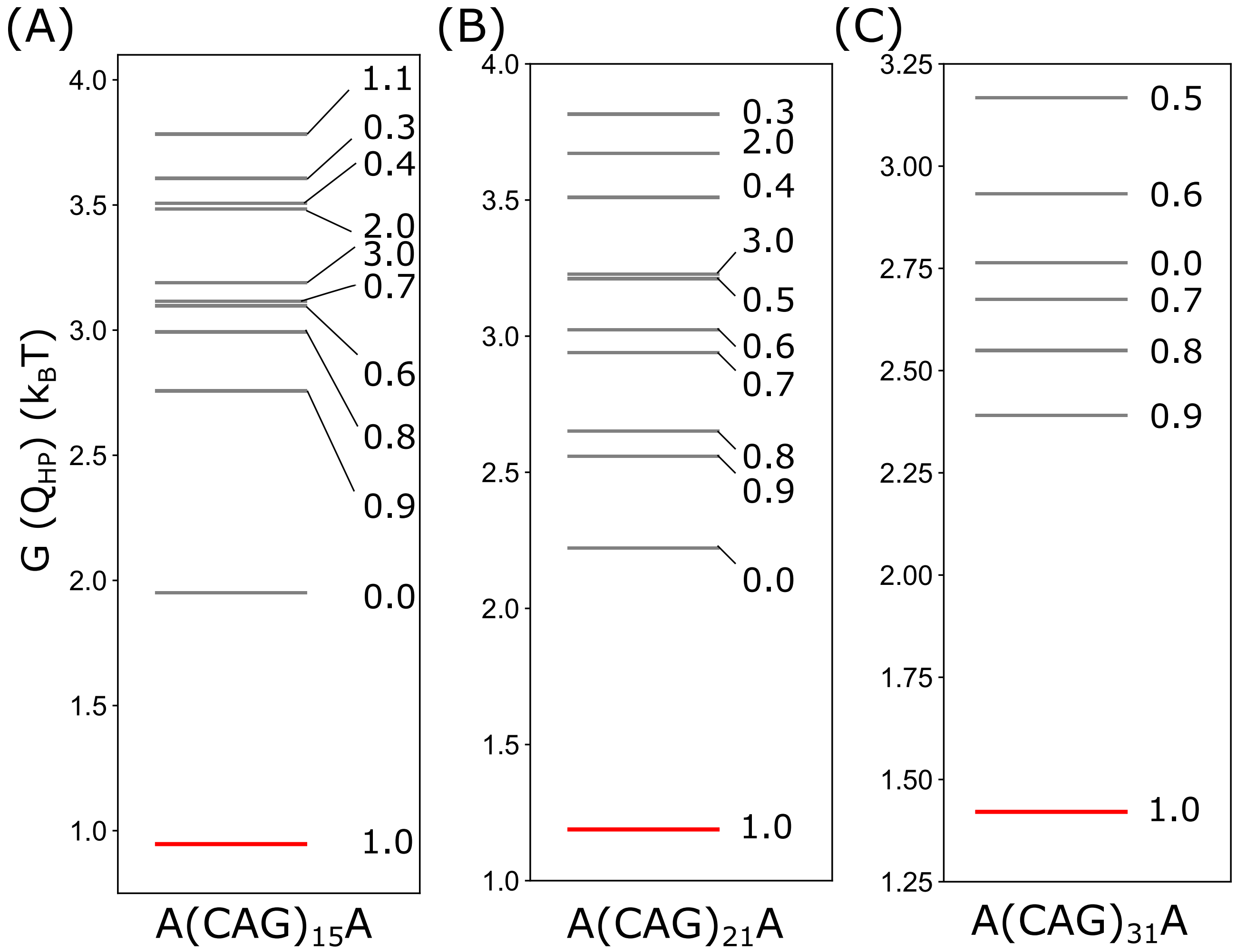


Figure S10: **Spectra of low free energy states $G(Q_{HP})$ for odd $n$ as a function of $Q_{HP}$.** (A) $G(Q_{HP})$ for odd numbered sequences as a function of $Q_{HP}$. (A) $G(Q_{HP})$ for A(CAG)$_{15}$A. (B) $G(Q_{HP})$ for A(CAG)$_{21}$A. (C) $G(Q_{HP})$ for A(CAG)$_{31}$A.

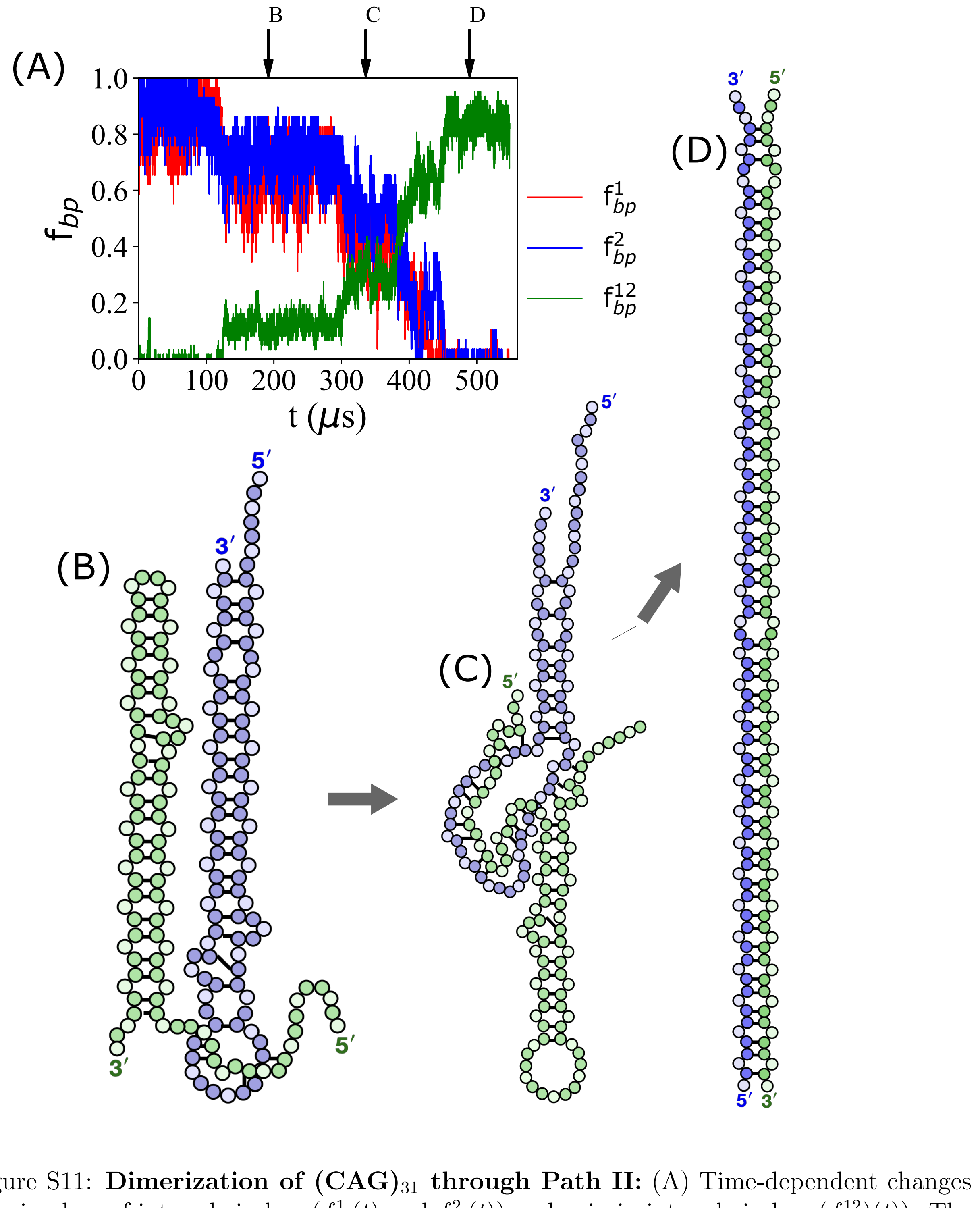


Figure S11: **Dimerization of $(\text{CAG})_{31}$ through Path II:** (A) Time-dependent changes showing loss of intra-chain bps ($f^1_{bp}(t)$ and $f^2_{bp}(t)$) and gain in inter-chain bps ($f^{12}_{bp})(t)$). The simulations were started with conformations in their ground states ($Q_{HP} = 1$). The value of $C_s$ is 100mM and $T$=37°C. (B and C) Initial hairpin conformations first transition to the structures which have a slipped ends at the termini. The hairpins with the slipped end start interacting with the loop region of the other chain and eventually lead to the formation of a dimer. Representative structures along the pathways are shown.

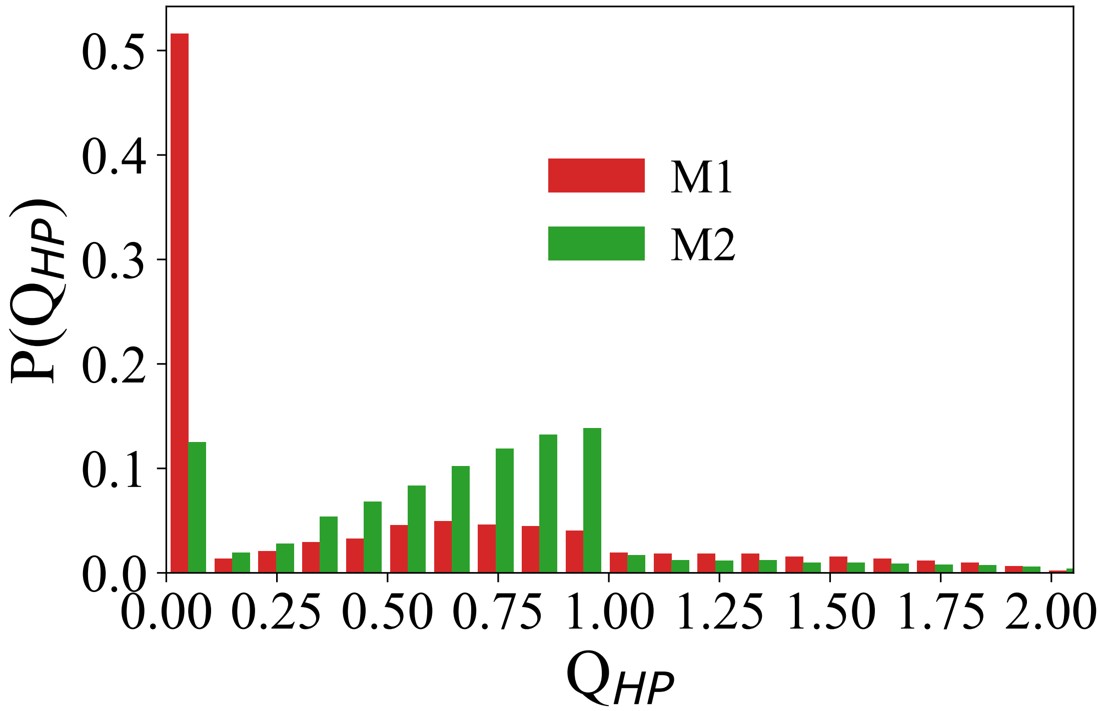


Figure S12: **Probability distribution of $Q_{HP}$ for the mutant sequence.** $P(Q_{HP}$ for M1 (A(CCG)(CAG)$_{28}$(CGG)A) and M2 (A(CCG)(CAG)$_{29}$(CGG)A) are in red and green, respectively. Replacement of A-A by C-G reduces the population in SH significantly in the mutant sequences with respect to the wild type sequences (A(CAG)$_{30}$A and A(CAG)$_{31}$A).

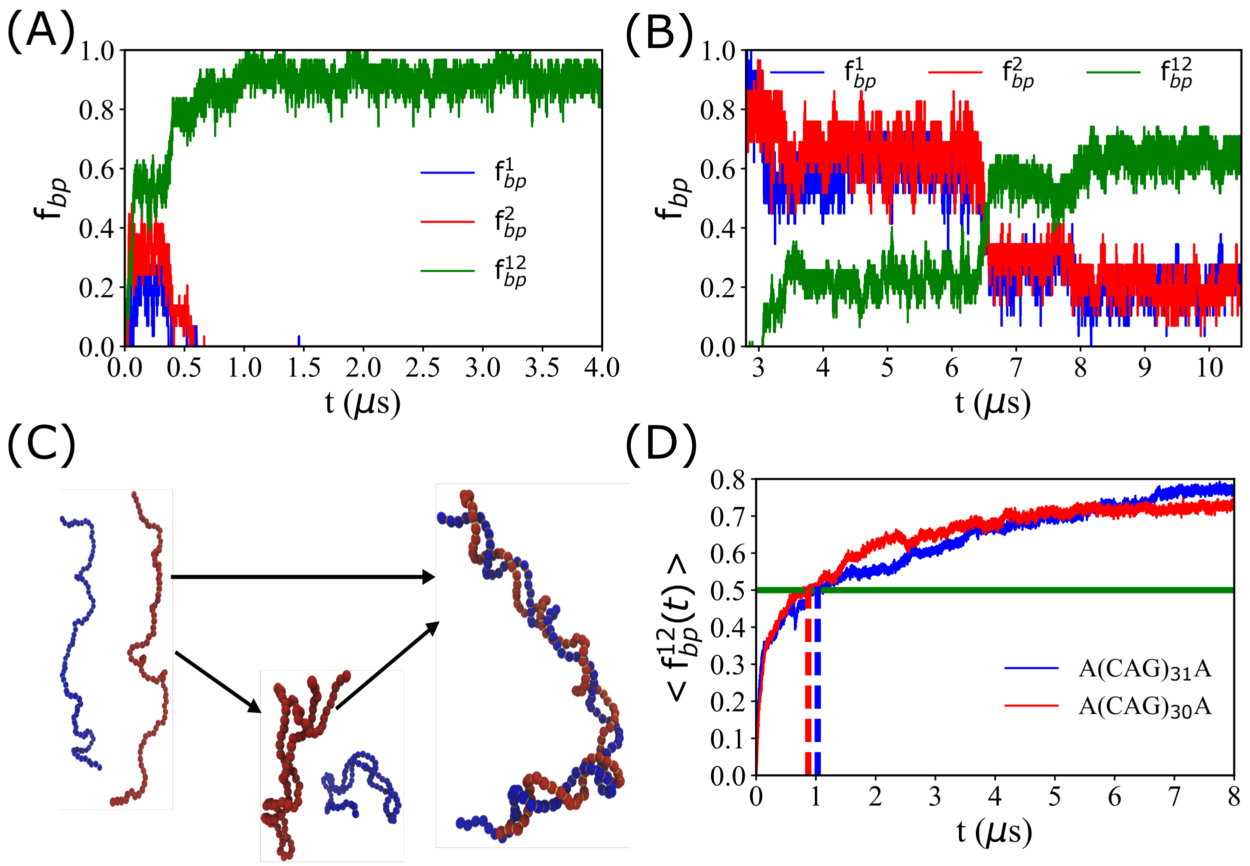


Figure S13: **Dimerization of A(CAG)$_{31}$A initiated from unfolded configurations.** (A) Fraction of intramolecular base pairs formed in chain 1 ($f^1_{bp}$), chain 2 ($f^2_{bp}$) and the fraction of intermolecular base pairs ($f^{12}_{bp}$) formed between two chains as a function of time are in blue, red and green, respectively. $f^1_{bp}$, $f^2_{bp}$ and $f^{12}_{bp}$ as a function of timed showed dimerization takes place directly from the unfolded state without forming significat intramolecular base pairs. (B) $f^1_{bp}$, $f^2_{bp}$ and $f^{12}_{bp}$ in red, blue and green indicates initially the monomers folded into the hairpin structures and with the advancement in simulations the hairpins start interacting with each other to form a dimer. (C) Schematic of the dimerization pathways started from the unfolded configurations. A dimer can form either from the monomers without populating the hairpin structures or with the formation of hairpins structures. (D) Time evaluation of intermolecualar base pairs, $\langle f^{12}_{bp}(t) \rangle$, for A(CAG)$_{31}$A (blue) and A(CAG)$_{30}$A (red) showed no significant difference in the rate of formation of dimer.

(A) M3 = A$(CCG)_3(CAG)_{25}(CGG)_3$A

M4 = A$(CAG)_{12}(CCG)_3$CAG$(CGG)_3(CAG)_{12}$A

(B)

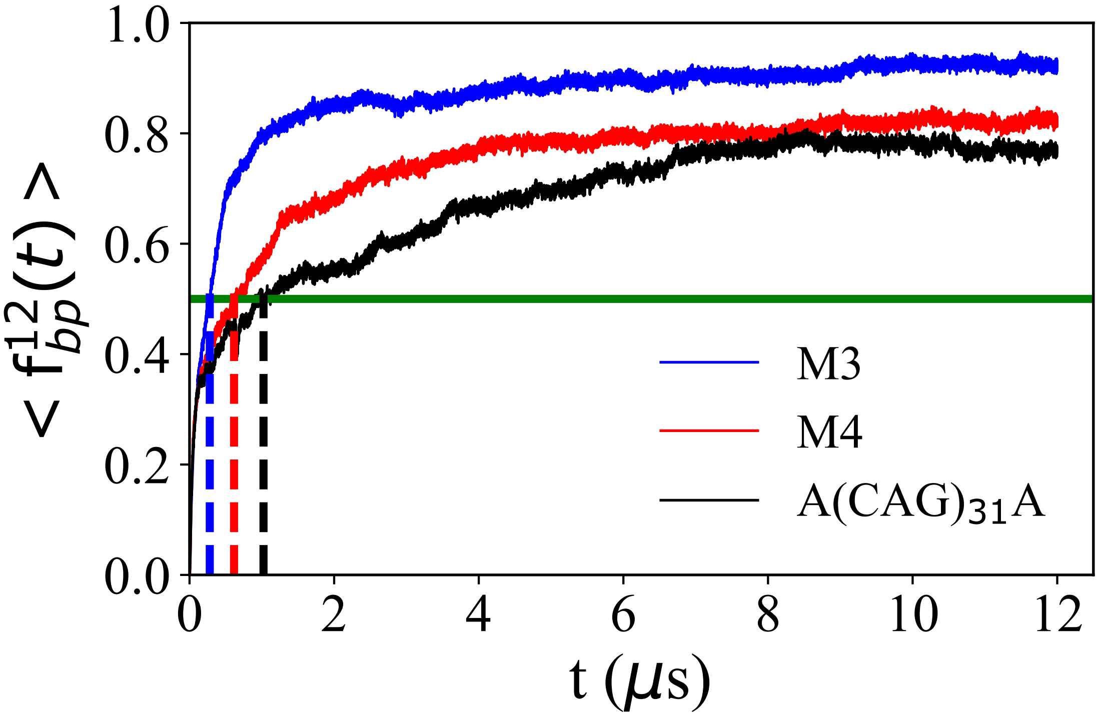


Figure S14: **Dimer formation in M3 and M4 mutants initiated from unfolded configurations.** (A) Sequences M3 and M4 used in the simulations. (B) Average fraction of intermolecular base pairs, $f_{bp}^{12}(t)$, formed between two chains as a function of time. Blue (red) is for the M3 (M4); $\langle f_{bp}^{12}(t) \rangle$ is calculated by averaging over 30 dimer forming trajectories. The dashed lines are qualitative estimates of dimerization time scale. The rate constant for dimerization of M3 is $\approx 2.2$ faster than M4.

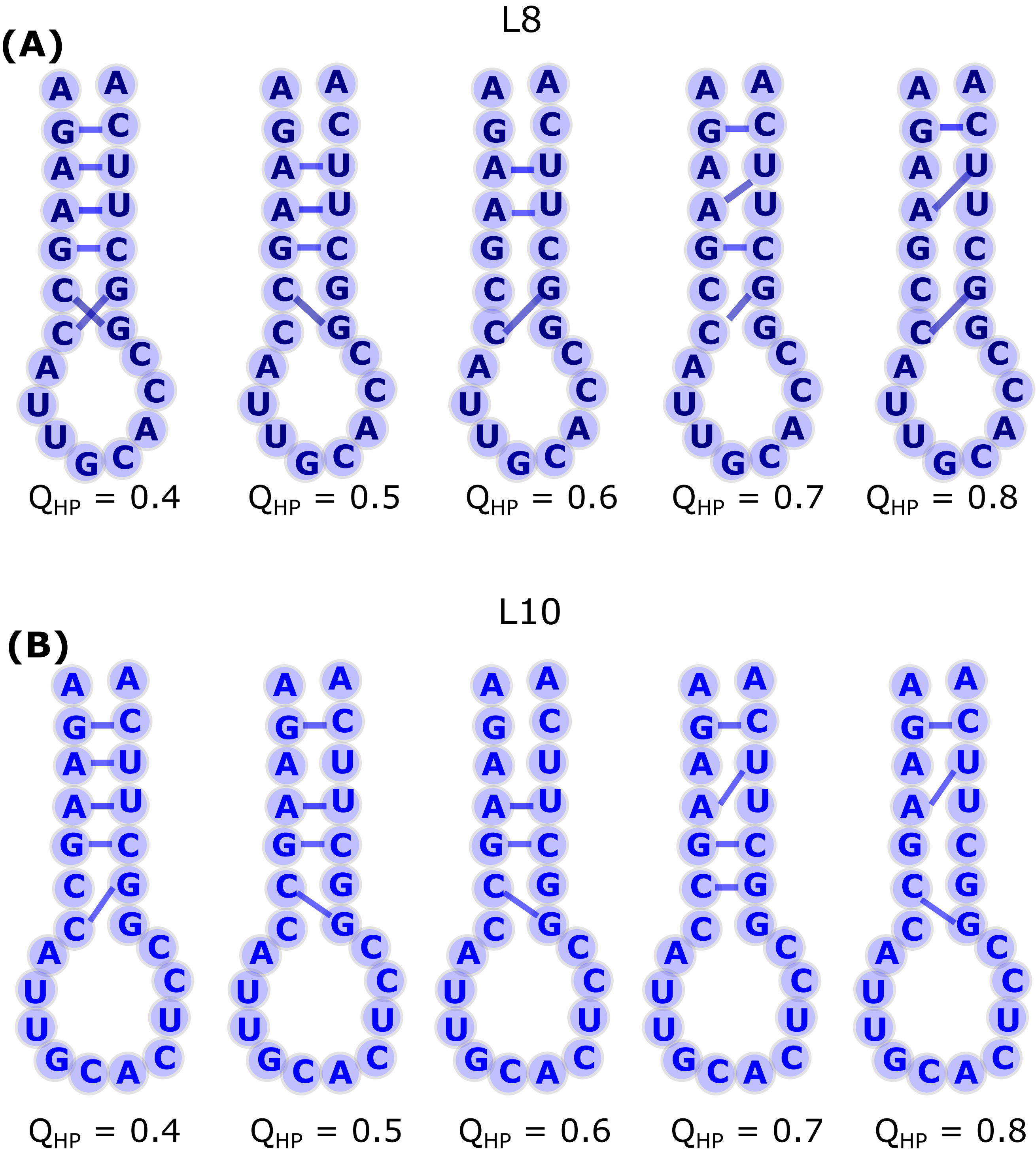


Figure S15: **Schematics of L8 and L10 configurations present between $Q_{HP} = 0$ and 1.7 in the free energy spectrum.** (A) Representative configurations of L8 corresponding to $Q_{HP} = 0.4$, 0.5, 0.6, 0.7 and 0.8 illustrates the $5'$ and the $3'$ are in close proximity. (B) Same as Figure A, except for sequence L10

### References


(1) Hyeon, C.; Dima, R. I.; Thirumalai, D. Pathways and kinetic barriers in mechanical unfolding and refolding of RNA and proteins. *Structure* **2006**, *14*, 1633–1645.

(2) Oosawa, F. *Polyelectrolytes*; New York, 1971.

(3) Manning, G. S. Limiting laws and counterion condensation in polyelectrolyte Solutions .I. colligative properties. *J. Chem. Phys.* **1969**, *51*, 924–933.

(4) Denesyuk, N. A.; Thirumalai, D. Coarse-Grained Model for Predicting RNA Folding Thermodynamics. *J. Phys. Chem. B* **2013**, *117*, 4901–4911.

(5) Malmberg, C.; Maryott, A. Dielectric constant of water from 0-degrees-c to 100-degree-c. *J. Res. Natl. Bur. Stand.* **1956**, *56*, 1–8.

(6) Dima, R.; Thirumalai, D. Exploring protein aggregation and self-propagation using lattice models: Phase diagram and kinetics. *Protein Sci.* **2002**, *11*, 1036–1049.

(7) Honeycutt, J. D.; Thirumalai, D. The Nature of Folded States of Globular-Proteins. *Biopolymers* **1992**, *32*, 695–709.

(8) Ermak, D.; McCammon, J. Brownian dynamics with hydrodynamic interactions. *J. Chem. Phys.* **1978**, *69*, 1352–1360.

(9) Kumar, S.; Bouzida, D.; Swendsen, R. H.; Kollman, P. A.; Rosenburg, J. The Weighted Histogram Analysis Method for Free-Energy Calculations on Biomolecules .1. The Method. *J. Comput. Chem.* **1992**, *13*, 1011–1021.

(10) Guo, Z.; Thirumalai, D. Kinetics of protein-folding - nucleation mechanism, time scales and pathways. *Biopolymers* **1995**, *36*, 83–102.

(11) Systematic generation and imaging of tandem repeats reveal base-pairing properties that promote RNA aggregation. *Cell Reports Methods* **2022**, *2*, 100334.

(12) Jain, A.; Vale, R. D. RNA phase transitions in repeat expansion disorders. *Nature* **2017**, *546*, 243+.
